# Analytical and functional clearance of dsRNA contaminants from *in vitro* transcribed mRNA by an engineered dsRNA-binding protein

**DOI:** 10.64898/2026.09.03.749252

**Authors:** Tyson Vonderfecht, Peter Lam, Genia Verovskaya, Jason Potter

**Author notes:** To whom correspondence should be addressed. Correspondence may also be addressed to.

## Abstract

Double-stranded RNA (dsRNA) is a biologically active contaminant of *in vitro* transcribed (IVT) mRNA that can reduce protein expression, induce cytokines, affect cell viability, and confound experimental results. Here, we show that dsRNA contaminants containing modified nucleotides are difficult to quantify and functionally assess. Common antibody-based dsRNA ELISAs under-detected pseudouridine- and N1-methylpseudouridine-modified dsRNA, while a higher-sensitivity ELISA improved detection but remained chemistry dependent. Defined dsRNA spike-in experiments showed that effects of dsRNA on cell viability, cytokine induction, and protein expression were chemistry dependent and occurred at low burdens of 0.05-0.5 ng dsRNA per µg RNA. Analytical dsRNA depletion across several workflows did not always predict suppression of cytokine induction, supporting the need for a functional cell-based readout. We engineered B2-S, a single-chain spliced variant of the Flock House virus B2 dsRNA-binding protein, to selectively deplete dsRNA from crude IVT reactions while preserving mRNA recovery and integrity. Among tested methods, B2-S was the only non-enzymatic, non-chromatographic workflow that reduced the cytokine marker IP-10/CXCL10 to mock-transfection levels. These findings establish a paired analytical and functional framework for IVT mRNA quality assessment and introduce B2-S as an accessible method for producing mRNA with improved potency and reduced immunostimulatory activity.

**GRAPHICAL ABSTRACT:** 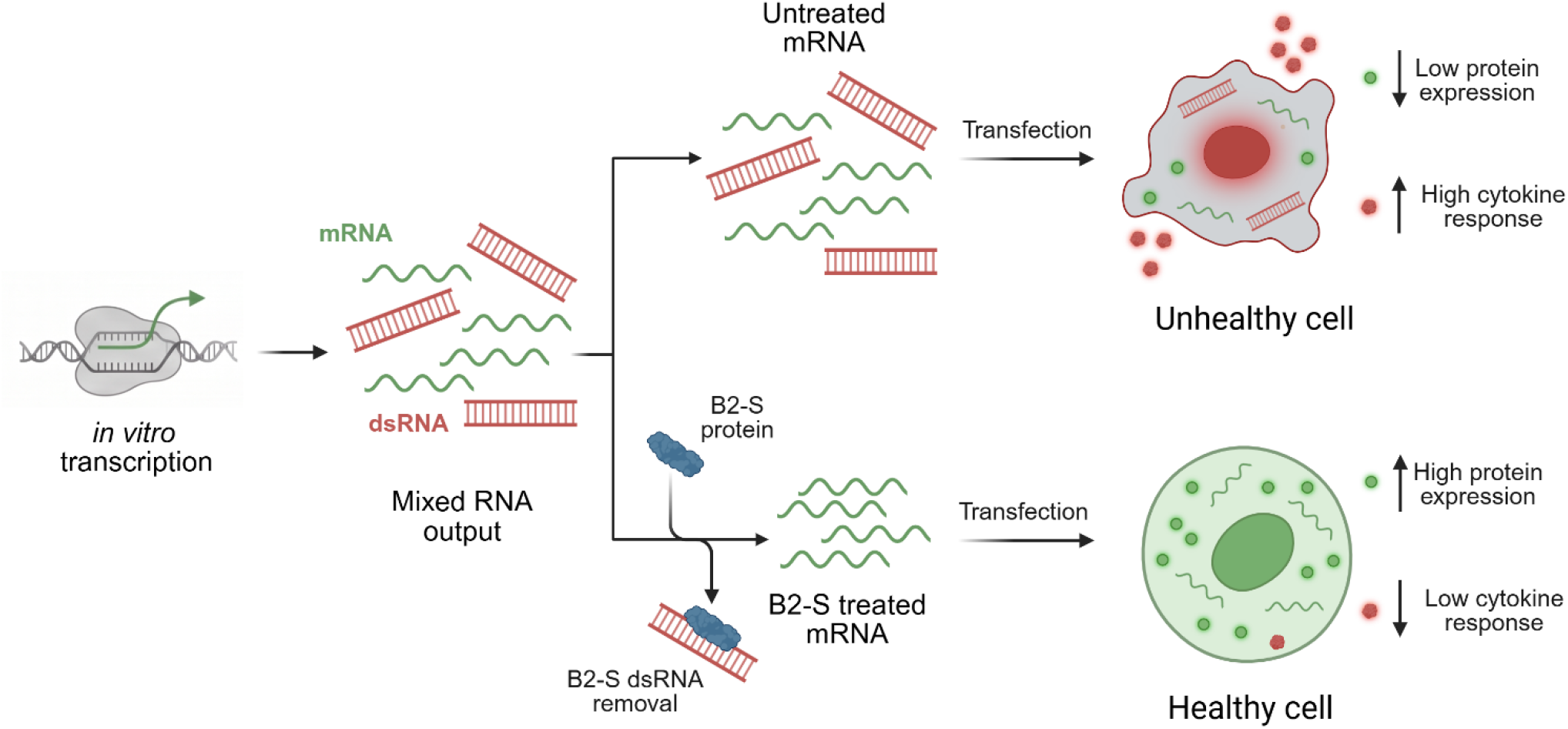

## INTRODUCTION

Messenger RNA (mRNA) is becoming a versatile and powerful tool across both therapeutic and research settings. It has emerged as a transformative platform in medicine for vaccines, protein replacement, and gene editing therapies. In research, mRNA is used for transient protein expression, genome editing, developmental biology studies, and somatic cell reprogramming, among other applications (1–8). Despite this promise, the advantage of mRNA is challenged by issues of immunogenicity and potency that are linked to the purity of the final mRNA product.

The mRNA synthesis process uses *in vitro* transcription (IVT), a cell-free process in which an RNA polymerase (RNAP) transcribes a linearized DNA template to create single-stranded RNA (ssRNA) (9). However, this reaction also produces unwanted biologically active byproducts, most notably double-stranded RNA (dsRNA) (10–13). These arise from aberrant RNA priming events within the reaction, where RNA fragments serve as primers for 3′ extension using RNA as a template to generate dsRNA species of both short and long lengths (Figure 1A) (12–14). The presence of dsRNA in IVT mRNA preparations poses a significant immunological challenge as mammalian cells contain receptors that detect dsRNA as part of antiviral defense, including endosomal Toll-like receptors, particularly TLR3, and cytosolic sensors such as RIG-I, MDA5, PKR, and the OAS/RNase L system (15,16). Activation of these pathways triggers type I interferon and pro-inflammatory cytokine responses, leading to global translational suppression, which reduces expression of the intended protein, as well as cellular toxicity and other inflammatory side effects (10–13,15,16). This response can limit therapeutic efficacy where robust and reproducible protein expression is required (10–13). Similar challenges occur in research settings, in which innate immune activation can confound experimental interpretation, reduce transfection efficiency, induce unwanted transcriptional and translational changes, and/or compromise cell viability (10–13). These challenges highlight the need for effective strategies to minimize or eliminate dsRNA contaminants in IVT mRNA preparations.

**Figure 1.**
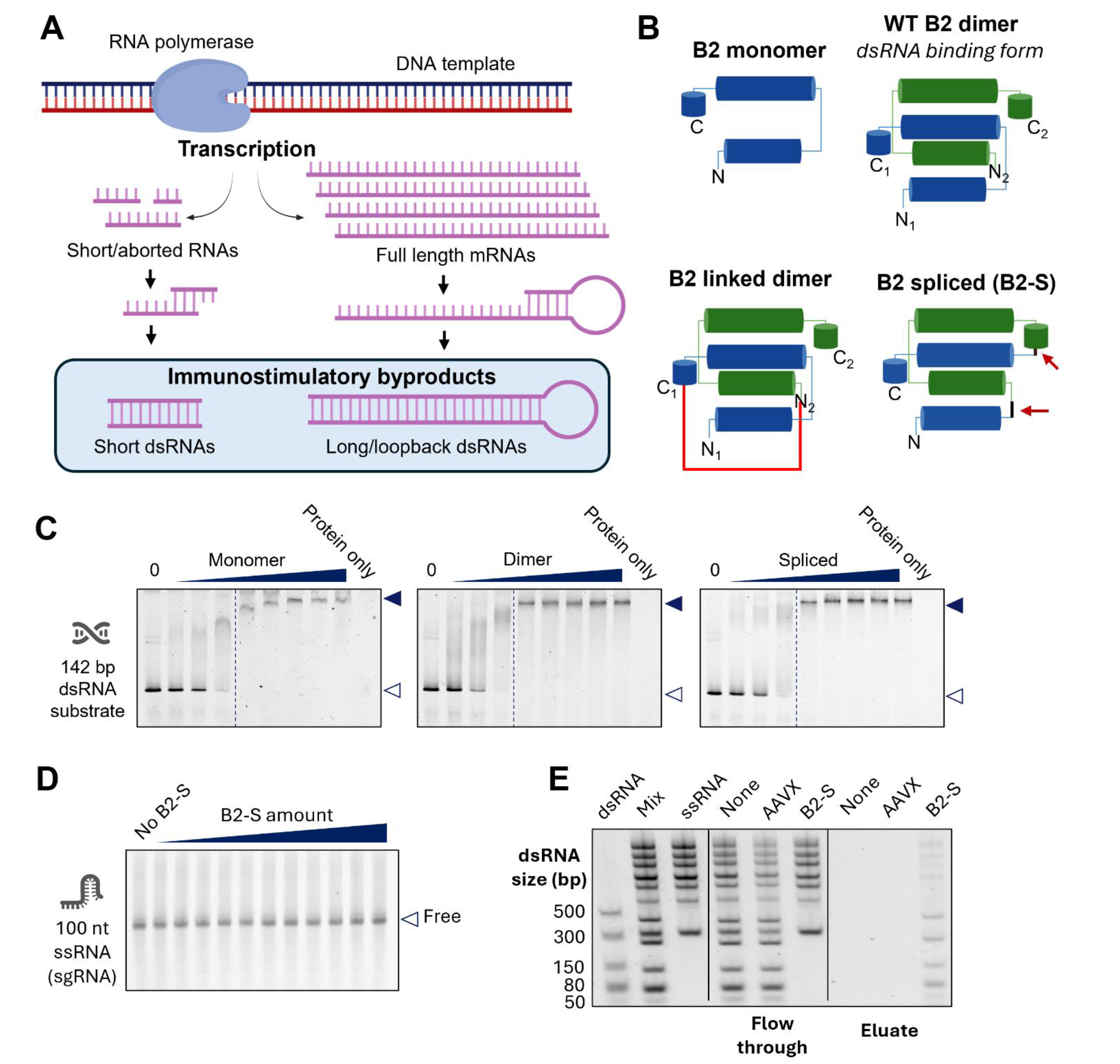
B2-S protein can remove dsRNA from ssRNA in mixed RNA samples. **(A)** Mechanism for dsRNA formation during IVT. DNA-dependent RNA polymerase generates short, aborted transcripts and full-length mRNAs. These RNAs can anneal to complementary sequences in *cis* or in *trans*, creating templates for RNA-dependent RNA synthesis by the polymerase and producing immunogenic short and long/loopback dsRNA byproducts (12–14). Image was made with BioRender. **(B)** Schematic of B2 protein architectures. Each cylinder represents an α-helix. Upper left, wild-type B2 monomer. Upper right, wild-type (WT) B2 dimer, the native dsRNA-binding unit, formed by two noncovalently associated subunits (subunit 1, blue; subunit 2, green). Bottom left, B2 linked dimer, generated by covalently joining two B2 monomers with a long flexible linker (red). Bottom right, B2 spliced single-chain construct, engineered by covalently joining the two subunits at the sites indicated by the red arrows to create a single polypeptide that mimics the native dimer conformation. Image was made with BioRender. **(C)** Electrophoretic mobility shift assays of B2 variants with a 142-bp dsRNA substrate. A constant amount of dsRNA substrate was incubated with increasing, molar-matched concentrations of monomer (left), dimer (middle), or spliced (right) B2 protein (lanes left to right). “Protein only” lanes contain protein alone with no dsRNA. “0” lanes contain dsRNA substrate alone with no protein. **(D)** A constant amount of 100-nucleotide CRISPR single-guide RNA (sgRNA), a structured ssRNA, was incubated with increasing amounts of B2-S protein (lanes left to right). “No B2-S” lane contains ssRNA alone with no protein. In (C) and (D), filled arrowheads indicate the bound protein–dsRNA complex and open arrowheads indicate free substrate. **(E)** dsRNA removal assay: “dsRNA” and “ssRNA” correspond to the respective ds- or ss-RNA ladders. “Mix” is the mixture of the two ladders and is the input sample. “None” is the input mix with no protein; “AAVX” is the input mix with a biotinylated anti-AAVX nanobody; and “B2-S” contains the input mix with B2-S protein, which carries a streptavidin-binding peptide (SBP) tag. Flow through is the sample after incubation with streptavidin beads, and eluate is the sample after biotin elution.

The dsRNA byproducts generated during IVT are not a single, well-defined impurity but a heterogeneous population with varying lengths and sequences, complicating their efficient removal during purification (12–14). Commonly used mRNA purification approaches, such as oligo(dT) capture, LiCl precipitation, or tangential flow filtration, are unable to selectively separate the dsRNA species from the desired mRNA product (17). As a result, several strategies have been developed to reduce or eliminate dsRNA contaminants from IVT mRNA preparations (17). These include reverse-phase high-performance liquid chromatography (RP-HPLC) (10), cellulose-ethanol purification (11), dsRNA-affinity chromatography (18), RNase III digestion (19), RNA polymerase engineering (12,13,20), or optimization of IVT conditions (21,22). These methods are limited by trade-offs in yield, mRNA integrity, scalability, operational complexity, or efficiency in removing dsRNA (17). Modified nucleotides such as N1-methylpseudouridine (m1Ψ) can reduce innate immune activation but do not eliminate activation driven by dsRNA byproducts (10,11,23,24). These challenges are further compounded by limitations in commonly used antibody-based methods for detecting and quantifying heterogeneous dsRNA contaminants (25,26). Furthermore, while dsRNA removal is known to improve IVT mRNA performance, the cellular effects of quantitatively defined dsRNA burdens in mRNA preparations have not been systematically established. It remains unclear how both dsRNA abundance and nucleotide chemistry influence innate immune activation and mRNA translation.

Here, we present B2-S, an engineered dsRNA-binding protein that enables selective affinity depletion of dsRNA from IVT mRNA preparations while preserving mRNA integrity and recovery. We define the dsRNA recognition properties of B2-S, develop a higher-sensitivity dsRNA ELISA for modified dsRNA detection, and show that ELISA alone is not sufficient to predict cellular innate immune activation. Using the IP-10/CXCL10 cytokine as an orthogonal biological readout, we demonstrate that B2-S removes the biologically active dsRNA burden across multiple IVT mRNAs, improving protein expression while reducing innate immune activation.

## MATERIAL AND METHODS

### General reagents

Ecosurf EH-9, Bond-Breaker TCEP solution, 100% Glycerol solution, 1M Tris, pH 7.5, DTT (No-Weigh Format), 2M triethylamine acetate, 0.5M EDTA, and 1M HEPES were all obtained from Thermo Fisher Scientific. Absolute ethanol, acetonitrile, NaCl, and bisTris were all purchased from Fisher Scientific.

### B2 design, cloning, and purification

The wild-type B2 protein sequence used in this study consists of amino acids 1 – 73 (UniProt ID P68831). The protein sequence for the linked dimer variant contained two B2 subunits with a 20-residue glycine linker in between. The spliced variant consisted of the first α-helix of one B2 subunit connected to a second B2 molecule via a GGSG linker and the remaining portion of the first subunit joined to the C-terminus of the second by another GGSG linker. All variants contained an N-terminal streptavidin binding peptide (SBP, see (27)) connected to the protein by a glycine linker.

DNA fragments encoding each B2 variant were synthesized by Thermo Fisher GeneArt™ services. The fragments were cloned by GeneArt Seamless Cloning and Assembly Kit (Thermo Fisher) into a modified pET151 expression vector (Thermo Fisher) where the sequence encoding the 6xHis-V5-TEV peptide was removed by PCR. All resulting expression plasmids were sequenced-verified and transformed into chemically competent *E. coli* BL21 Star (DE3) cells (Thermo Fisher).

Expression was performed in cells grown in Terrific Broth medium and induced overnight with IPTG (Thermo Fisher) at 20°C. Cells expressing the proteins were pelleted by centrifugation, resuspended in a bisTris-NaCl buffer with EcoSurf EH-9, and lysed by high-pressure homogenization (Avestin Emulsiflex C5). Tagged proteins were purified at 4°C on an AKTA Pure FPLC instrument by running the lysate through a column with High-Capacity Streptavidin Agarose Resin (Thermo Fisher) and eluted with 10 mM biotin. Proteins were further purified by anion exchange chromatography (POROS™ HQ resin, Thermo Fisher), and then cation exchange chromatography (POROS XS resin, Thermo Fisher). Purified proteins were dialyzed into a low salt Tris-TCEP buffer with 50% glycerol. Purity of eluted proteins was assessed by SimplyBlue™ SafeStain (Thermo Fisher) staining of protein separated on denaturing 4-12% NuPAGE™ Bis-Tris Mini Protein Gels (Thermo Fisher). Gels were imaged on an iBright™ FL1500 Imaging System (Thermo Fisher). Purification yields were estimated from absorbance at 280 nm based on extinction coefficients computed from protein amino acid composition. Purified proteins were aliquoted and stored at −20°C.

### B2 binding assays

The complex between B2 and dsRNA or sgRNA substrates was monitored by electrophoretic mobility shift assays (EMSA). 50 ng of a 142-bp dsRNA substrate (Jena Bioscience) or 35 ng of sgRNA (Thermo Fisher TrueGuide™ Synthetic gRNA; spacer sequence: GGGAAAGACCCAGCAUCCGU) was equilibrated with a series of two-fold serial dilutions of B2 protein (12 µM – 47 nM) in B2 binding buffer (Tris-NaCl-EDTA-TCEP) at room temperature for at least 30 minutes. Samples were mixed 1:1 with 1:10 diluted TBE Hi-Density Sample Buffer (Thermo Fisher). Protein–RNA complexes were resolved from unbound RNA by nondenaturing PAGE in Novex™ TBE Gels, 6% (Thermo Fisher) at room temperature. Gels were removed from the cassettes, stained with 1X SYBR™ Gold (Thermo Fisher) in water for 5 minutes, washed in water for 1 minute, and then imaged on an iBright instrument.

The ssRNA and dsRNA ladder binding assays were performed by mixing 500 ng RiboRuler™ High Range RNA Ladder (Thermo Fisher) with 250 ng dsRNA ladder (New England Biolabs) in B2 binding buffer. 5 µL of B2-S or CaptureSelect™ Biotin Anti-AAVX Conjugate (Thermo Fisher) was added. Solutions were incubated at room temperature for 5 minutes and then transferred to microtubes containing 100 µL Dynabeads™ Streptavidin Magnetic Bead slurry (Thermo Fisher) washed twice with B2 binding buffer. After a 5-minute incubation, a magnet (DynaMag™-2 Magnet, Thermo Fisher) was applied, and the supernatant (flow through) was transferred to a clean microtube. A 10 mM biotin solution was used to resuspend the magnetic beads and elute the protein from the beads. After a 5-minute incubation, the magnet was applied and the supernatant (eluate) was transferred to a clean microtube. The RNA in the flow through and eluant samples were recovered by the GeneJET™ RNA Cleanup and Concentration Micro Kit (Thermo Fisher) and eluted into 15 µL water. 12 µL of sample was mixed with 3 µL E-Gel™ sample loading buffer (Thermo Fisher) and loaded onto a 2% E-Gel 48 agarose gel. The gel was imaged on an iBright instrument.

### mRNA synthesis and purification

DNA templates encoding GFP, RFP, EPO, or FLuc all contained a T7 RNA polymerase promoter with an AG initiation for use with CleanCap™ AG, followed by the human alpha or beta globin 5’ untranslated region (UTR) with a Kozak sequence (CACC) and then the open reading frame (ORF). Following the ORF was the human alpha or beta globin 3’ UTR. These sequences were ordered as plasmids from GeneArt and then PCR amplified by SuperFi™ II Master Mix (Thermo Fisher) with a reverse primer containing a 120 nt poly(T) stretch (Integrated DNA Technologies (IDT)) to create a DNA amplicon with the T7 promoter, UTRs, ORF, and encoded poly(A) tract. PCR conditions recommended by the kit user guide were used. The PCR product size was confirmed by gel electrophoresis, purified by PureLink™ PCR Purification kit (Thermo Fisher), and diluted to 100 ng/µL in water.

The purified DNA amplicon was then used for mRNA synthesis by IVT with the mMESSAGE mMACHINE™ T7 mRNA Kit with CleanCap Reagent AG (Thermo Fisher) by following the manufacturer’s instructions. Modified mRNAs were made by complete substitution of uridine with Ψ, m1Ψ, or 5moU (all from Thermo Fisher). All mRNAs were purified with the Dynabeads RNA purification kit (Thermo Fisher) by following the manufacturer’s instructions and eluted into THE RNA Storage Solution (sodium citrate buffer, Thermo Fisher). Concentration and purity (A260/A280 and A260/A230) were measured using a NanoDrop™ spectrophotometer (Thermo Fisher).

mRNA synthesis with a low dsRNA mutant T7 RNA polymerase was performed using the Codex HiCap RNA T7 Polymerase (Aldevron) (20) by following the manufacturer’s instructions except a 0.8:1 CleanCap:NTP ratio was used to help ensure high capping efficiency. The NTPs and cap analog used were from the mMESSAGE mMACHINE T7 mRNA Kit with CleanCap Reagent AG. Inorganic pyrophosphatase (Thermo Fisher) and RiboLock™ RNase Inhibitor (Thermo Fisher) were both added per the manufacturer’s instructions. mRNA purification was performed as described previously.

The low dsRNA optimized IVT condition was performed using urea as described in Piao et al. (21). This reaction was set up using the mMESSAGE mMACHINE T7 mRNA Kit with CleanCap Reagent AG, and urea (Thermo Fisher) was added to a final concentration of 1M. mRNA purification was performed as described previously.

mRNA integrity was assayed by capillary electrophoresis using a TapeStation instrument (Agilent) with RNA ScreenTape and Sample Buffer (Agilent) by following the manufacturer’s instructions. Integrity for some samples was also assayed by denaturing agarose gel electrophoresis using a 1% E-Gel (Thermo Fisher). 1 µL of 200 ng/µL mRNA was mixed with 9 µL RNA Gel Loading Dye (Thermo Fisher), heated at 70°C for 10 minutes, and then chilled on ice for 5 minutes. 8 µL of the mixture was loaded onto the E-Gel. The RiboRuler High Range RNA Ladder (Thermo Fisher) was used for RNA sizing. Gels were imaged by an iBright instrument.

### dsRNA removal by B2-S

dsRNA removal from the IVT mRNA by B2-S was performed by diluting the RNA sample 5- fold in B2 binding buffer and adding 5 µL of B2-S protein. Samples were incubated at room temperature for 5 minutes and then transferred to microtubes containing 100 µL washed streptavidin magnetic beads. After a 5-minute room temperature incubation, the tubes were transferred to a 37°C heat block and incubated for another 5 minutes to minimize nonspecific interactions with the magnetic beads. A magnet was then applied, and the supernatant was transferred to a clean microtube. The mRNA was purified from the supernatant using the Dynabeads RNA purification kit as described previously.

### dsRNA removal by HPLC

dsRNA was removed from purified mRNA samples as described previously (28). Briefly, 100 µg purified mRNA was loaded onto a 1260 Infinity HPLC system (Agilent Technologies) equipped with a Clarity™ 5 µm Oligo-RP 150 x 4.6 mm column (Phenomenex) set to 65°C. A linear gradient of buffer B (0.1 M triethylammonium acetate, pH 7.0, 25% acetonitrile) from 38% to 70% in buffer A (0.1 M triethylammonium acetate, pH 7.0) over 15 min at 1 mL/min was applied. Fractions were collected using an Agilent Technologies 1260 Infinity II Fraction Collector and concentrated with Thermo Scientific™ Pierce™ Protein Concentrators (PES, 30K MWCO). The mRNA was purified and exchanged into sodium citrate buffer using the Dynabeads RNA purification kit as described previously.

### dsRNA removal by cellulose

dsRNA removal by cellulose-ethanol purification was performed by exactly following the protocol as described in Baiersdörfer et al. (11). The mRNA was recovered and concentrated using the Dynabeads RNA purification kit as described previously.

### dsRNA removal by RNase III

dsRNA removal by RNase III was performed using the CellScript Min-Immune Gold dsRNA Removal Kit by following the manufacturer’s instructions. The mRNA was recovered and concentrated using the Dynabeads RNA purification kit as described previously.

### dsRNA removal by Vendor affinity resin

The Repligen AVIPure dsRNA Clear OPUS loose resin was used as the commercially available affinity resin (18). The protocol was optimized to identify the conditions for effective dsRNA removal and mRNA recovery. The loose resin was prepared by transferring 30 µL of stock resin slurry to a microcentrifuge fitted with a cellulose acetate spin column (Pierce Spin Cups - Cellulose Acetate Filter, Thermo Fisher). The resin slurry was centrifuged at 1000xg for 2 minutes and washed twice with buffer containing 50 mM HEPES, pH 7.2, 2 mM EDTA, and 700 mM NaCl. 100 µg of purified mRNA was diluted to 400 ng/µL in the HEPES buffer and then mixed well with the washed resin in the spin column. After a 10-minute room temperature incubation with shaking, the sample was spun down at 1000xg for 2 minutes. The mRNA was recovered from the flow through and concentrated using the Dynabeads RNA purification kit as described previously.

### dsRNA synthesis

The forward RNA strand was synthesized using the control GFP DNA template in the mMESSAGE mMACHINE T7 mRNA Kit with CleanCap Reagent AG to create a 0.9 kb RNA product that was either unmodified or fully substituted with Ψ, m1Ψ, or 5moU. The DNA template encoding the reverse strand was created by PCR with the control GFP DNA template and primers removing the original T7 promoter in the forward direction and adding a T7 promoter in the reverse direction (see Supplemental Figure 2A and Supplemental Methods). The reverse RNA strand was synthesized using this DNA template with the mMESSAGE mMACHINE T7 mRNA Kit with CleanCap Reagent AG to create a 0.9 kb RNA product that was either unmodified or fully substituted with Ψ, m1Ψ, or 5moU. Both RNA products were purified by LiCl precipitation as described in the mMESSAGE mMACHINE T7 mRNA Kit with CleanCap Reagent AG user guide. The dsRNA byproduct was removed by cellulose-ethanol purification as described previously. The two strands were annealed by mixing equimolar amounts of each in annealing buffer (10 mM Tris pH 7.5, 20 mM NaCl, and 1 mM EDTA) and incubating at 75°C in a heat block. After 5 minutes, the heat block was turned off to allow the sample to cool to room temperature (2-3 hours). The sample was purified by the GeneJET RNA Cleanup and Concentration Micro Kit and eluted into water. The concentration was measured by NanoDrop™ spectrophotometer using the dsDNA setting. Gel electrophoresis and dsRNA dot blot were performed to confirm dsRNA synthesis (see Supplemental Figures 2B and C).

### dsRNA dot blots

2 µL of IVT mRNA (200 ng/µL) was added to a Biodyne B Pre-Cut Modified Nylon Membrane (0.45 µm, Thermo Fisher) so that each dot corresponds to 400 ng mRNA. The dsRNA standard consisted of 2 µL of 0.5 ng/µL m1Ψ dsRNA (described previously). The samples were added by multichannel pipette, making sure the tips did not touch the membrane. The membrane was allowed to dry overnight at room temperature. On the next day, the membrane was blocked for 2–3 hours on a shaker at room temperature in a solution containing 2 mL WesternBreeze Diluent™ A and 3 mL WesternBreeze Diluent B (Thermo Fisher) with 5 mL water. The membrane was rinsed twice with water, and the primary antibody solution (7 mL water, 2 mL Diluent A, and 1 mL Diluent B mixed with 2 µL J2 anti-dsRNA antibody (1:5000 dilution; Thermo Fisher)) was added. The membrane in the primary antibody solution was incubated on a shaker at room temperature overnight. The solution was removed, and the membrane was washed 5 times in 20 mL PBST (PBS + 0.1% Tween-20 (Thermo Fisher)). The secondary antibody solution (7 mL water, 2 mL Diluent A, and 1 mL Diluent B mixed with 2 µL Invitrogen™ Goat anti-Mouse IgG2a Secondary Antibody, HRP (1:5000 dilution; Thermo Fisher)) was added. The membrane in the secondary antibody solution was incubated on a shaker at room temperature for 2 hours. The solution was removed, and the membrane was washed 5 times in 20 mL PBST. The membrane was rinsed twice with water and the SuperSignal™ West Pico PLUS Chemiluminescent Substrate (Thermo Fisher) was added according to the manufacturer’s protocol. After 5 minutes, the membrane was imaged on an iBright imaging system.

The dot blots evaluating the sensitivity of different anti-dsRNA antibodies to chemically-modified dsRNA (Figure 2C) were prepared by serially diluting the samples 1:2 and adding 2 µL to the membranes. The blots were probed for dsRNA using the J2 antibody (Thermo Fisher), K1 antibody (Thermo Fisher), K2 antibody (Exalpha Biologicals), or 9D5 antibody (Absolute Antibody). J2 and K1 were detected by the Invitrogen Goat anti-Mouse IgG2a Secondary Antibody as described previously. K2 was detected by the Invitrogen Goat anti-Mouse IgM Secondary Antibody, HRP (Thermo Fisher), and 9D5 by the Invitrogen Goat anti-Mouse IgG1 Secondary Antibody, HRP (Thermo Fisher). All secondary antibodies were used at 1:5000 dilutions. Blots were visualized as described previously.

**Figure 2.**
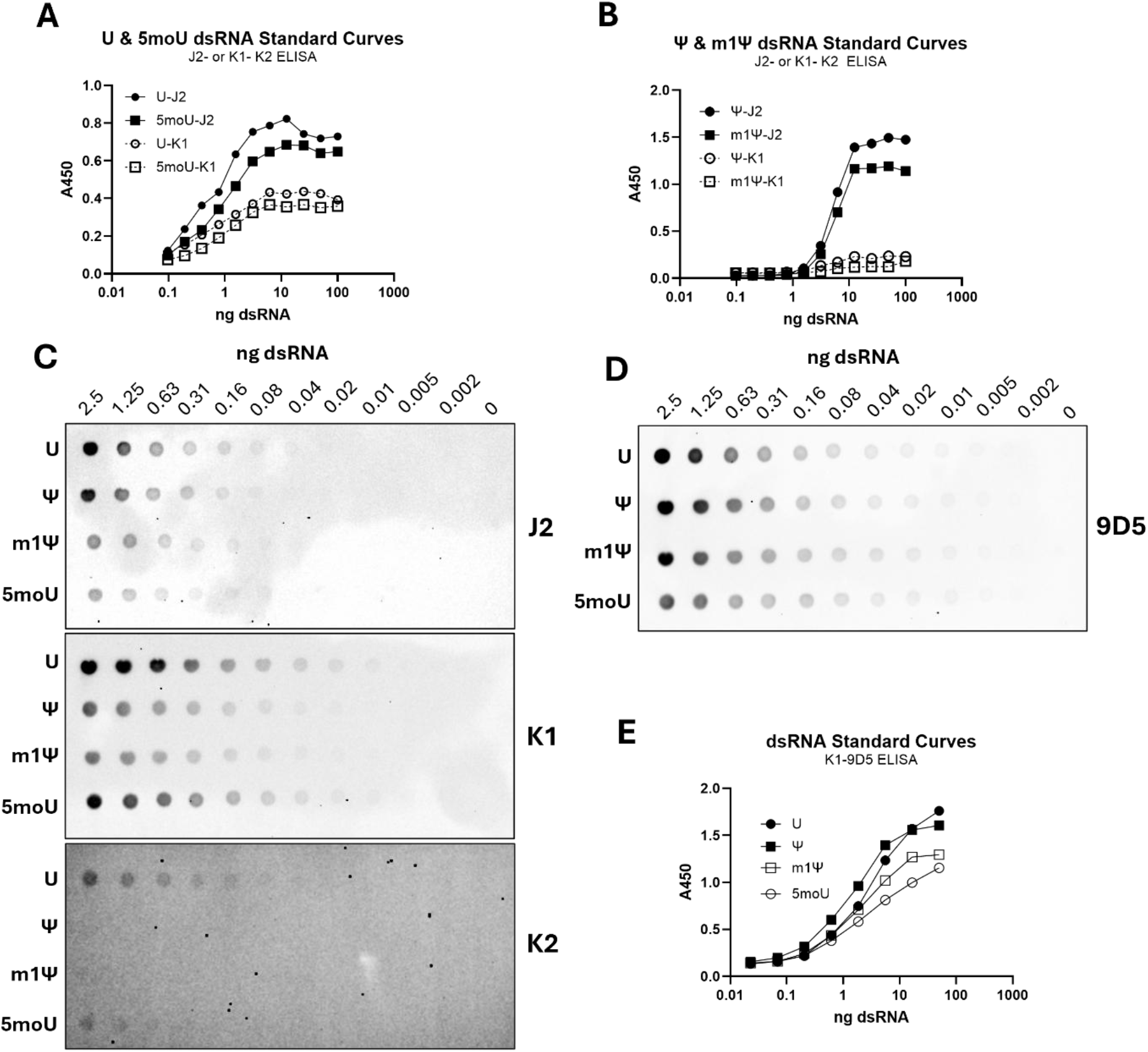
Chemically-modified dsRNA affects immunoassay sensitivity. **(A)** Standard curves generated with a ∼1-kb dsRNA containing U or 5moU from a commercial dsRNA ELISA with the commonly used J2-K2 or K1-K2 antibody pairs. **(B)** The same antibody pairs demonstrate reduced sensitivity for Ψ- or m1Ψ-modified dsRNA. **(C)** Serial dilutions of ∼1- kb dsRNA containing U, Ψ, m1Ψ, or 5moU were spotted onto membranes and probed with the anti-dsRNA antibodies J2 (top), K1 (middle), or K2 (bottom). **(D)** Serial dilutions of the same dsRNA species in (C) were probed with the anti-dsRNA antibody 9D5. **(E)** Standard curves generated with ∼1-kb dsRNA containing U, Ψ, m1Ψ, or 5moU using K1 as the capture antibody and 9D5 as the detection antibody in a sandwich ELISA.

### dsRNA ELISA

Standard curves of unmodified or fully substituted dsRNA with Ψ, m1Ψ, or 5moU from J2-K2 or K1-K2 ELISA systems were created using the commercially available J2-K2 or K1-K2 SCICONS anti-dsRNA ELISA kits (Exalpha Biologicals) by following the manufacturer’s recommended protocol.

The K1-9D5 ELISA system was performed by coating white ELISA plates (Thermo Fisher) with a solution of the K1 anti-dsRNA mouse IgG2a antibody diluted in PBS, pH 7.4 (Thermo Fisher). After an overnight incubation at 4°C, the plates were blocked with SuperBlock™ Blocking Buffer (PBS, Thermo Fisher) for 1 hour at room temperature. The dsRNA standards were prepared from a 10 pg/µL stock by performing 1:2 dilutions into assay buffer (SuperBlock Blocking Buffer diluted 1:10 into PBS). 50 ng of RNA in 100 µL assay buffer were added to the plates and incubated on a shaker for 1 hour at room temperature. Plates were washed 4 times with PBST. Detection antibody solution (9D5 anti-dsRNA mouse IgG1 antibody diluted into assay buffer) was added to the plates and incubated on a shaker for 1 hour at room temperature. Plates were washed 4 times with PBST. Secondary antibody solution (Goat anti-Mouse IgG1 Secondary Antibody, HRP, diluted into assay buffer) was added to the plates and incubated on a shaker for 1 hour at room temperature. Plates were washed 5 times with PBST, and development began by adding TMB-ELISA Substrate solution (Thermo Fisher). Development was stopped after ∼5 minutes by adding Stop Solution (Thermo Fisher).

Absorbance at 450 nm was read on a Varioskan™ LUX Microplate Reader (Thermo Fisher). All standard curves were constructed using Prism (GraphPad), and best-fit equations were generated in Prism using a 4PL sigmoid fit with the unknowns being interpolated from the standard curve. Each standard or sample was assayed in duplicate.

### Cell culture and transfection

The JAWSII murine immature dendritic and BJ human fibroblast cell lines were obtained from the American Type Culture Collection (ATCC) and cultured according to ATCC recommendations. All media and supplements were obtained from Thermo Fisher.

20,000 BJ or 40,000 JAWSII cells were seeded in 100 µL media onto 96-well flat bottom cell culture plates (Thermo Fisher) for mRNA transfections. BJ cells were transfected with 200 ng EPO mRNA by Lipofectamine™ MessengerMAX™ Transfection Reagent (Thermo Fisher) by following the manufacturer’s 0.3-µL protocol. BJ cells were transfected with 250 ng RFP mRNA by Lipofectamine 2000 Transfection Reagent (Thermo Fisher) by following the manufacturer’s 0.5-µL protocol. The corresponding amounts of RFP mRNA in Figures 3F-H were transfected into JAWSII cells using MessengerMAX reagent as described previously.

**Figure 3.**
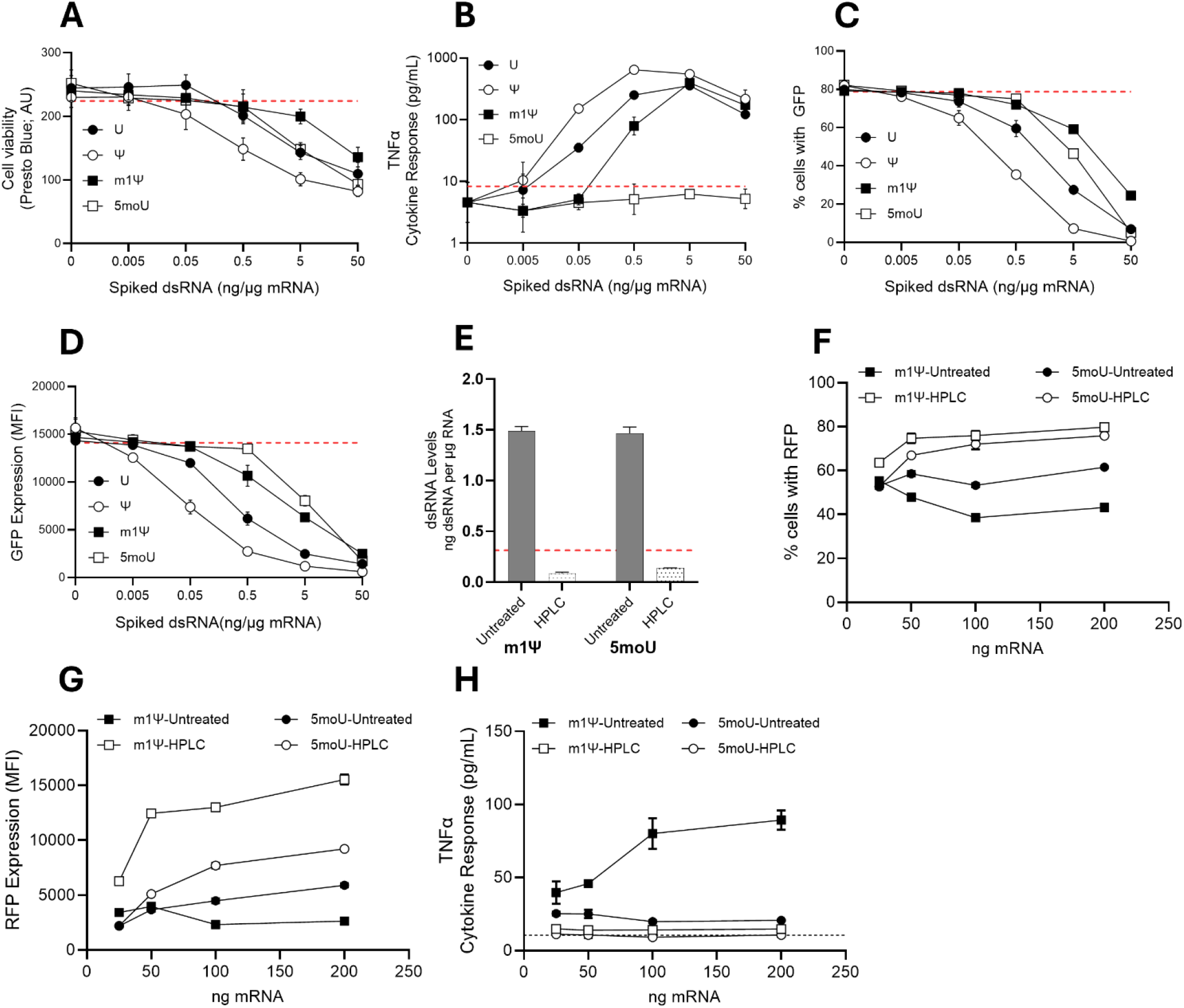
dsRNA burdens impair mRNA performance in a dose- and chemistry- dependent manner. (A-D) JAWSII murine immature dendritic cells transfected with HPLC- purified GFP mRNA (minimal dsRNA) and increasing amounts of spiked-in dsRNA. Cell viability (A), TNFα secretion (B), the percentage of cells expressing GFP (C), and GFP expression (D) were measured. The red dotted lines represent the lower or upper standard deviation of the GFP only transfection control. Error bars represent standard deviations. N=3. **(E)** dsRNA levels in m1Ψ or 5moU RFP mRNAs as measured by K1-9D5 ELISA. **(F-H)** After transfecting JAWSII cells with the m1Ψ or 5moU RFP mRNAs, the percentage of cells expressing RFP (F), RFP expression (G), and TNFα secretion (H) were measured. The red dotted line in (E) represents the LLOQ of the dsRNA ELISA (0.3125 ng dsRNA per µg RNA) as described in the text. The dotted line in (H) represents the background TNFα levels as measured in a mock transfection control. Error bars represent standard deviations. N=3.

Samples for the dsRNA spike-in experiments were performed by preparing a 50 ng/µL HPLC-purified 5moU GFP mRNA in sodium citrate buffer and mixing in dsRNA at 2.5 ng/µL. The mixture was serially diluted 1:10 into the 50 ng/µL HPLC-purified 5moU GFP mRNA solution. 2 µL of each dilution was transfected into JAWSII cells using MessengerMAX reagent as described previously. At least three transfections were performed for all conditions. Each transfection experiment was performed at least two times.

### Cell assays and data analysis

RFP or GFP analysis of BJ or JAWSII cells was performed by flow cytometry using an Attune CytPix Cytometer. Live cells were stained with CellTrace Calcein Violet AM (Thermo Fisher) by following the manufacturer’s instructions. The following gating strategy was performed: 1) SSC-A vs FSC-A to gate on cells and remove debris; 2) SSC-A vs SSC-H to gate on single cells; 3) FSC-H vs VL1 to gate on live cells; and then FSC-H vs BL1-A or YL1-A to gate on GFP or RFP cells, respectively.

The levels of TNFα in 50 µL JAWSII cell media were measured with the Invitrogen Mouse TNF alpha Uncoated ELISA Kit (Thermo Fisher) by following the manufacturer’s instructions. The levels of IP-10/CXCL10 in 15 µL BJ cell media were measured with the Invitrogen/PeproTech Human IP-10 (CXCL10) ELISA Development Kit (TMB) by following the manufacturer’s instructions. Absorbances were read by the Varioskan microplate reader. All standard curves were constructed as described previously.

EPO levels in 1:20 diluted BJ cell media were measured with the Invitrogen Human EPO ProQuantum Immunoassay Kit (Thermo Fisher) by following the manufacturer’s instructions. The EPO assay was run on a QuantStudio 7 Flex Real-Time PCR System (Thermo Fisher) and data analysis was performed using the online application software.

Cell viability was measured with the PrestoBlue™ Cell Viability Reagent (Thermo Fisher) by following the manufacturer’s instructions. Fluorescence was read on the Varioskan microplate reader.

BJ cells were prepared for staining by washing them twice with dPBS (Thermo Fisher), fixing them with 10% formalin (Scigen) for 10 minutes, and permeabilizing them with PBST for 5 minutes. Cells were washed with twice with dPBS and blocked in BlockAid™ Blocking Solution (Thermo Fisher) for 30 minutes. Cells were stained for 30 minutes in blocking buffer containing Alexa Fluor™ Plus 555 Phalloidin (1:400 dilution, Thermo Fisher) and DAPI (1 µg/mL, Thermo Fisher). Following 4 washes with dPBS, the cells were visualized on an EVOS™ M5000 Imaging System.

All statistical analyses were performed using Prism (GraphPad). The statistical tests used for individual experiments are specified in the corresponding figure legends. The *in vivo* total flux values were background-adjusted and log-transformed prior to statistical analysis. Specifically, values were transformed as log10(total flux + c), where c was defined as the mean total flux of the PBS control animals. PBS was included as a negative control for visualization but was excluded from the primary factorial analysis. The groups were analyzed by two-way ANOVA. Comparisons between purification methods were performed within each timepoint using Bonferroni’s multiple comparisons test.

### mRNA-LNP formulation

FLuc mRNA was encapsulated using Vivofectamine™ VF232 Liver LNP composition in Ethanol (Thermo Fisher) following the manufacturer’s recommendations. mRNA was diluted in 30 mM sodium acetate buffer pH 5.5 (Thermo Fisher) to 0.21 mg/mL and complexed with VF232 Liver at a flow rate of 12 mL/min and ratio 3:1 using a microfluidic instrument (NanoAssemblr Ignite, Precision NanoSystems). The resulting mRNA-LNP was dialyzed in PBS pH 7.4 for 4 hours using Float-A-Lyzer Dialysis Device (Spectrum). Particle size was analyzed using dynamic light scattering (ZetaSizer Advance Pro Malvern Panalytical), and encapsulation efficiency was measured using Quant-iT™ RiboGreen™ Assay (Thermo Fisher) and analyzed on the Varioskan microplate reader (Thermo Fisher).

### Animal experiments

All animal experiments were approved by Institutional Animal Care and Use Committee. 1 mg/kg FLuc mRNA-LNPs were delivered to 8-week-old female BALB/c (Jackson Laboratory) mice using tail vein injections. Control mice were injected with 200 µL PBS. Four or 24 hours after mRNA-LNP injection, the mice were injected intraperitoneally with 100 µL of IVISbrite™ D-Luciferin BIOluminescent Substrate in RediJect™ Solution (PerkinElmer). Mice were anesthetized using isoflurane, and the luciferase signal was analyzed *in vivo* and *ex vivo* using an IVIS™ Lumina LT Series III In Vivo Imaging System (Revvity). Serum was collected by retroorbital bleeding using micro-hematocrit capillary tubes (Fisherbrand™) into 0.8 mL serum separator tubes (MiniCollect) and then centrifuged at 2500xg for 15 minutes. Cytokine analysis was performed using ProcartaPlex™ Mouse Cytokine & Chemokine Panel 1A, 36- plex (Thermo Fisher) and a Luminex FLEXMAP3D instrument system (Thermo Fisher) by following the manufacturer’s recommended protocol. Cytokine data visualization was performed using a Python 3 notebook in JupyterLab (Anaconda). For analysis of liver biochemistry, serum samples were sent to Antech Diagnostics.

AI tools were used for editorial assistance and language refinement. All scientific content, interpretations, and final text were reviewed and approved by the authors.

## RESULTS

### Engineered B2-S selectively binds dsRNA and separates dsRNA from ssRNA mixtures

The dsRNA-binding protein B2 from Flock House virus was selected as a scaffold for developing a dsRNA removal system because of its high specificity for dsRNA that is independent of its nucleotide sequence (29–31). B2 recognizes dsRNA through a dimerization-dependent mechanism (Figure 1B) (29,30), prompting us to design three variants for evaluation: (1) a monomeric form, (2) a covalently linked dimer in which two monomers were connected by a long flexible linker, and (3) a “spliced” single-chain variant engineered to form a stable dimeric structure. In the spliced variant, the first α-helix of one B2 subunit is connected to a second B2 molecule via a short linker, and the remaining portion of the first subunit is joined to the C-terminus of the second by another short linker, producing a single protein that mimics the dimer conformation (Figure 1B). Each construct was expressed in *E. coli*, column-purified, and evaluated for specificity toward a 142-bp dsRNA substrate by electrophoretic mobility shift assays (EMSA). All three proteins shifted the substrate over a similar protein-concentration range, indicating comparable binding to this dsRNA substrate (Figure 1C).

We selected the B2-spliced variant (B2-S) for further characterization because its single-chain design was expected to simplify binding kinetics and support downstream use as a reagent for removing dsRNA contaminants from IVT mRNA preparations. To test ssRNA binding, we performed EMSA binding assays using a 100-nucleotide CRISPR single-guide RNA (sgRNA), a structured ssRNA containing three stem-loop motifs with dsRNA-like character (Supplemental Figure 8A) (32). We chose the sgRNA as a model structured ssRNA because mRNAs commonly form intramolecular secondary structures, including hairpins containing short base-paired stem regions (33), raising the possibility that a dsRNA-binding protein could also recognize structured regions within the intended mRNA product. B2-S produced no detectable band shift and did not deplete the free sgRNA signal, indicating minimal detectable binding to this structured ssRNA substrate (Figure 1D).

The minimum dsRNA-binding length of B2-S was estimated to be 21 bp, although weak binding to a 19-bp dsRNA substrate was observed (Supplemental Figure 1A). This slightly differs from the wild-type B2 monomer, which weakly recognized the 17-bp dsRNA substrate (Supplemental Figure 1B). These results somewhat contrast a previous report showing that wild-type B2 can bind 17-, 19-, and 21-bp dsRNA substrates with similar affinities (29). This discrepancy may be due to different assay conditions and/or dsRNA substrates.

We then tested 21-bp dsRNA substrates containing internal mismatches (Supplemental Figure 1C). One- or two-base mismatches substantially weakened binding, indicating that B2-S prefers fully base-paired dsRNA. In contrast, an internal G-U wobble pair did not measurably reduce binding, consistent with its preservation of the A-form helix (34).

We next tested whether B2-S could selectively separate dsRNA from ssRNA in mixed RNA populations. To enable affinity-based capture, the streptavidin-binding peptide (SBP) tag at the N-terminus of B2-S was used as a purification handle (Supplemental Figure 1D). B2-S was incubated with a mixture of ssRNA and dsRNA ladders and then captured by streptavidin magnetic beads. After bead capture, biotin was added to competitively elute the B2-S protein, along with any bound RNA substrate. RNA in all samples was recovered by silica spin column purification and analyzed by agarose gel electrophoresis to determine the distribution of ssRNA and dsRNA. A biotinylated anti-AAV nanobody (AAVX) was included as a non-dsRNA binding control to assess nonspecific capture. As shown in Figure 1E, B2-S efficiently depleted dsRNA from the flow-through, leaving ssRNA predominantly unbound, and enriched dsRNA in the bound fraction. A faint signal from the ssRNA ladder was observed in the B2-S-bound fraction, indicating either trace carryover or low-level recognition of structured or dsRNA species within the ladder. In contrast, the AAVX control did not deplete dsRNA from the flow-through, with both ssRNA and dsRNA remaining in the unbound fraction. These results demonstrate that B2-S can selectively separate dsRNA from ssRNA in mixed RNA samples.

### Modified dsRNA species are differentially detected in antibody-based assays

Residual dsRNA quantification is necessary for IVT mRNA quality assessment. The commonly used J2 antibody-based dot blot assay is useful for relative comparisons but is only semi-quantitative and does not provide reliable absolute quantification (25,26). Sandwich enzyme-linked immunosorbent assays (ELISAs) or other immunoassays using dsRNA-specific antibodies provide a quantitative alternative, with several commercial kits now available. However, their performance depends on antibody recognition of dsRNA, which can be affected by nucleotide modification (25,26). Therefore, we evaluated whether commonly used J2-K2 or K1–K2 ELISA formats (35) could sensitively and reproducibly quantify unmodified and modified dsRNA species during development of the B2-S dsRNA removal system.

Both ELISA formats were evaluated using a ∼1 kb dsRNA species that was either unmodified (U) or fully substituted with pseudouridine (Ψ), N1-methylpseudouridine (m1Ψ), or 5- methoxyuridine (5moU) (see Supplemental Figures 2A-C). For U and 5moU dsRNA, both J2- K2 and K1–K2 ELISAs demonstrated broad dynamic range and strong signal intensity (Figure 2A). In contrast, both formats showed reduced signal intensity and compressed dynamic range for Ψ- and m1Ψ-modified dsRNA (Figure 2B). These results indicate that J2-K2 and K1- K2 ELISA platforms under-detect Ψ- and m1Ψ-modified dsRNA and may underestimate dsRNA levels in IVT mRNA preparations with these modifications.

To better understand the sensitivities of the J2, K1, and K2 toward modified dsRNA species, we performed dot blot analyses with these antibodies (Figure 2C). Both J2 and K1 detected all dsRNA species across a broad range of amounts with K1 exhibiting slightly higher sensitivity than J2. K2 produced lower overall signal, a narrower detection range, and markedly weaker detection of Ψ- and m1Ψ-modified dsRNA relative to U and 5moU dsRNA. These results suggest that the reduced performance of the J2–K2 and K1–K2 ELISA formats with Ψ- and m1Ψ-modified dsRNA is likely driven by limited recognition by the K2 detection antibody rather than inefficient capture by J2 or K1. The modification-dependent antibody responses also show that dsRNA standards should match the nucleotide chemistry of the IVT mRNA sample for absolute quantification.

Next, we screened several dsRNA-specific antibodies by dot blot to identify a stronger detection antibody for a K1-capture ELISA. Among the antibodies tested, 9D5 demonstrated robust detection of all dsRNA species across a broad range (Figure 2D). We therefore paired K1 capture with 9D5 detection (K1-9D5 ELISA) and evaluated performance across the modified dsRNA panel (Figure 2E). This antibody pairing produced strong signals and broad dynamic range for all tested dsRNA chemistries. The improvement was most evident for Ψ- and m1Ψ-modified dsRNA, where K1-9D5 expanded the standard-curve range and improved low-input signal discrimination. The assay still showed chemistry-dependent responses, reinforcing the need for modification-matched standards for absolute quantification.

The lower limit of quantification (LLOQ) for K1-9D5 was established using both spike-in recovery and standard curve performance. HPLC-purified m1Ψ-modified RFP or Cas9 mRNA was supplemented with known amounts of m1Ψ-modified dsRNA and analyzed by ELISA. Measured concentrations matched expected values down to 0.3125 ng dsRNA per µg RNA (Supplemental Figure 3A). At lower spike-in levels, calculated dsRNA concentrations were increasingly affected by assay background, reducing confidence in quantification. The J2-K2 ELISA produced no detectable signal for the same spiked samples, further showing that K2- based detection underestimates m1Ψ-modified dsRNA (data not shown).

Standard curves from three independent ELISA runs showed that 0.3125 ng dsRNA per µg RNA marked the transition from the low-signal area of the curve to the more responsive portion of the assay, with limited signal separation below 0.3125 ng/µg mRNA and a sharper increase in A450 above this point (Supplemental Figure 3B). Based on this and the spike-in recovery data, we define 0.3125 ng dsRNA per µg RNA as the practical LLOQ for the K1-9D5 ELISA. Below this concentration, background increasingly contributed to the calculated dsRNA concentration and limited reliable quantification.

### Defined dsRNA burdens impair IVT mRNA performance in a chemistry-dependent manner

With a quantitative assay in place, we asked if the assay’s LLOQ was biologically relevant and how defined dsRNA burdens affect IVT mRNA performance in cells. Although dsRNA impurities are widely recognized as detrimental, many studies have evaluated their biological effects by comparing IVT mRNA preparations before and after dsRNA reduction or removal procedures (10–13,18–22). Such comparisons do not directly establish the cellular response to a defined dsRNA burden in mRNA background, nor whether that response depends on dsRNA nucleotide chemistry. To address this, we spiked HPLC-purified 5moU-modified GFP mRNA (containing minimal dsRNA) with increasing amounts of defined dsRNA species containing U, Ψ, m1Ψ, or 5moU. The preparations were transfected into JAWSII immature dendritic cells, a sensitive innate immune model with strong dsRNA-sensor activity (36). Cell viability was assayed to assess toxicity, TNFα secretion was measured as a marker of innate immune activation (36), and GFP-positive cells and GFP expression intensity were quantified by flow cytometry to evaluate mRNA function (Figures 3A-D).

Increasing dsRNA burden impaired mRNA performance in a dose- and chemistry-dependent manner. Ψ dsRNA reduced cell viability beginning near 0.5 ng dsRNA per µg mRNA. U, m1Ψ, and 5moU dsRNA reduced viability at higher spike levels (Figure 3A). TNFα secretion was also chemistry dependent: U, Ψ, and m1Ψ dsRNA induced TNFα at low spike levels (0.05-0.5 ng/µg), whereas 5moU dsRNA produced little or no TNFα response across the tested range (Figure 3B). Ψ and U dsRNA reduced both the percentage of GFP-positive cells and GFP intensity at 0.5 ng/µg. m1Ψ dsRNA produced intermediate effects, and 5moU dsRNA was least disruptive (Figures 3C and 3D). Overall, the disruptive potency of dsRNA species followed the trend: Ψ > U > m1Ψ > 5moU.

Functional impairment occurred at dsRNA levels as low as 0.05-0.5 ng/µg for most chemistries. The K1-9D5 LLOQ of 0.3125 ng dsRNA per µg RNA falls within this biologically active range, suggesting that the LLOQ is biologically relevant. We therefore used 0.3125 ng/µg as a working analytical target for subsequent experiments, not as a universal biological specification.

Because 5moU-modified dsRNA was apparently less disruptive than m1Ψ-modified dsRNA in the spike-in study, we next tested whether 5moU offered an advantage at the mRNA- payload level. We generated m1Ψ- or 5moU-modified RFP mRNA by IVT and used HPLC purification to reduce dsRNA below the working threshold (Figure 3E). Different amounts of untreated (IVT mRNA purified by LiCl precipitation without HPLC purification) or HPLC-purified versions of each mRNA were transfected into JAWSII cells. After 24 hours, we measured RFP-positive cells, RFP intensity, and TNFα secretion (Figures 3F-H).

HPLC purification improved both modified mRNAs, confirming that dsRNA removal improves IVT mRNA activity (10). However, purified m1Ψ-modified mRNA produced higher RFP expression than purified 5moU-modified mRNA across the tested dose range (Figure 3G). This higher expression did not produce a marked TNFα increase after HPLC purification (Figure 3H). A similar trend was observed with GFP mRNA, additional constructs, and other cell systems (Supplemental Figure 4). These data support the use of m1Ψ modification combined with dsRNA removal to achieve optimal mRNA performance with minimal toxicity.

### B2-S removes biologically active dsRNA impurities from IVT mRNA

We next tested whether B2-S could reduce dsRNA below the working analytical target directly from IVT material. We generated m1Ψ-modified EPO mRNA, applied B2-S affinity capture directly to the crude IVT reaction, and then performed standard RNA purification, which in this case was RNA purification with Dynabeads (see Materials and Methods). We compared recovery, integrity, residual dsRNA, and biological activity against mRNA prepared by the standard purification alone (the untreated condition).

B2-S treatment preserved mRNA yields, with less than 10% yield loss, and did not affect integrity (Figures 4A and 4B). Notably, only B2-S-treated mRNA had dsRNA levels below the K1-9D5 ELISA LLOQ of 0.3125 ng dsRNA per µg mRNA, whereas untreated mRNA remained well above this level (Figure 4C). The dsRNA dot blot supports this observation (Figure 4D). Similar performance across mRNA constructs ranging from 0.8 kb to 5 kb further supports the broader applicability of the B2-S approach for dsRNA removal from diverse IVT mRNA products (Supplemental Figure 5).

**Figure 4.**
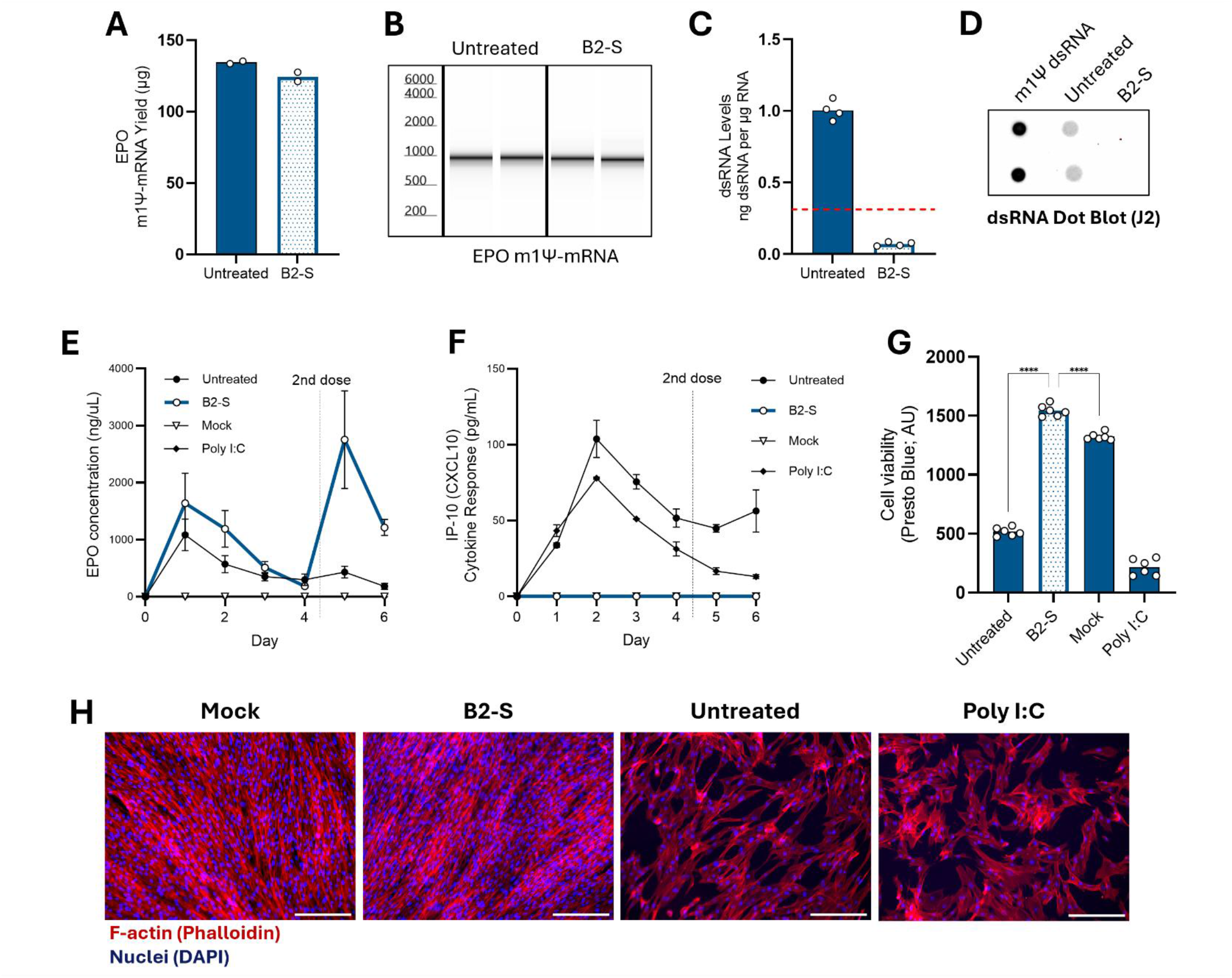
dsRNA depletion by B2-S improves EPO mRNA function. (A-D) EPO mRNA yields (A), integrity (B), and dsRNA levels as measured by K1-9D5 ELISA (C) or detected by dot blot (D). The red dotted line in (C) represents the ELISA LLOQ as described in the text. **(E and F)** EPO levels (E) and IP-10/CXCL10 secretion (F) in BJ fibroblasts after transfection with untreated or B2-S treated EPO mRNA. The vertical dotted line indicates when the cells were re-transfected with EPO mRNA. Bars represent the standard deviations. N=6. **(G)** Cell viability of the cells at the end of the experiment as measured by PrestoBlue cell viability reagent. Bars represent the mean, and individual replicate values are shown as open circles. Statistical significance was determined in (G) by one-way ANOVA followed by Dunnett’s multiple comparisons test versus the B2-S sample. Comparisons that are not significant are not indicated. **\*\*\*\***: P≤0.0001. **(H)** Representative fluorescence images of F-actin stained with phalloidin (red) and nuclei stained with DAPI (blue) at the end of the experiment. Scale bars, 300 µm. Mock is a transfection control with only the delivery reagent (no mRNA). Poly(I:C) served as a dsRNA-positive control.

We next evaluated whether dsRNA depletion by B2-S improved cellular performance. Untreated or B2-S-treated m1Ψ-modified EPO mRNA was transfected into human BJ fibroblasts, a primary-like human cell model suitable for assessing mRNA-associated cytotoxicity and immune stimulation (35). Media were collected and replaced daily for four days. Cells were then re-transfected, and media collection continued for an additional two days. Secreted EPO protein and the interferon-inducible chemokine IP-10/CXCL10 were measured in the conditioned media. IP-10/CXCL10 was selected as the primary cytokine readout because it is a sensitive marker of the antiviral interferon response induced by dsRNA-sensing pathways, providing a functional measure of any residual immunostimulatory activity after purification (35,37). At the end of the experiment, cell viability was measured, and cells were fixed and stained for F-actin and nuclei to assess cellular morphology.

B2-S-treated EPO mRNA had higher EPO expression than untreated EPO mRNA for the first two days. After that, the two conditions had similar EPO levels (Figure 4E) until the second dose. Re-transfection with untreated mRNA did not restore expression, whereas B2-S-treated mRNA significantly increased expression (Figure 4E). Thus, B2-S treatment improved expression durability and supported productive repeat transfection.

Untreated EPO mRNA induced IP-10/CXCL10, while B2-S-treated mRNA kept IP-10/CXCL10 at or near the baseline level (Figure 4F). Untreated mRNA also reduced the viability assay signal, whereas B2-S-treated mRNA preserved signal near or slightly above mock-transfected levels (Figure 4G). The modestly higher PrestoBlue signal for B2-S may reflect greater metabolic activity rather than higher cell number, potentially due to earlier contact inhibition in mock cells (38). Morphology matched the cell viability readouts: B2-S-treated mRNA retained organized F-actin and nuclear staining comparable to the mock transfection control, while untreated mRNA caused disruption like poly (I:C), a dsRNA mimic (10) (Figure 4H).

Collectively, these results show that B2-S reduces dsRNA contamination in IVT mRNA to below the level associated with cellular IP-10/CXCL10 response while preserving mRNA translation, viability, and morphology. Similar improvements in EPO expression and IP- 10/CXCL10 suppression were observed in additional cell lines (Supplemental Figure 6), supporting the broader utility of B2-S treatment across cellular contexts.

### B2-S compares favorably with tested dsRNA-reduction workflows while preserving mRNA recovery and integrity

We next compared B2-S with IVT-based dsRNA-reduction strategies and post-IVT dsRNA- removal methods to test whether analytical dsRNA depletion and functional clearance are equivalent (Figure 5A). For IVT reaction-based conditions, m1Ψ-modified RFP mRNA was generated using either a low-dsRNA-producing mutant T7 RNA polymerase or using urea supplementation during the IVT reaction (20,21), followed by standard RNA purification. For post-IVT conditions, m1Ψ-modified RFP mRNA was generated using wild-type T7 RNA polymerase and subjected to standard RNA purification, followed by dsRNA removal using HPLC (10), cellulose (11), RNase III treatment (19), or a commercially available dsRNA-affinity resin (Vendor resin) (18), and a final standard purification step. B2-S capture was evaluated in two configurations: applied directly to the crude IVT reaction mixture prior to standard RNA purification or applied after an initial standard purification followed by a final standard purification step. The mRNA yield, integrity, residual dsRNA levels, and functional performance were assessed across all conditions (Figures 5B–F).

**Figure 5.**
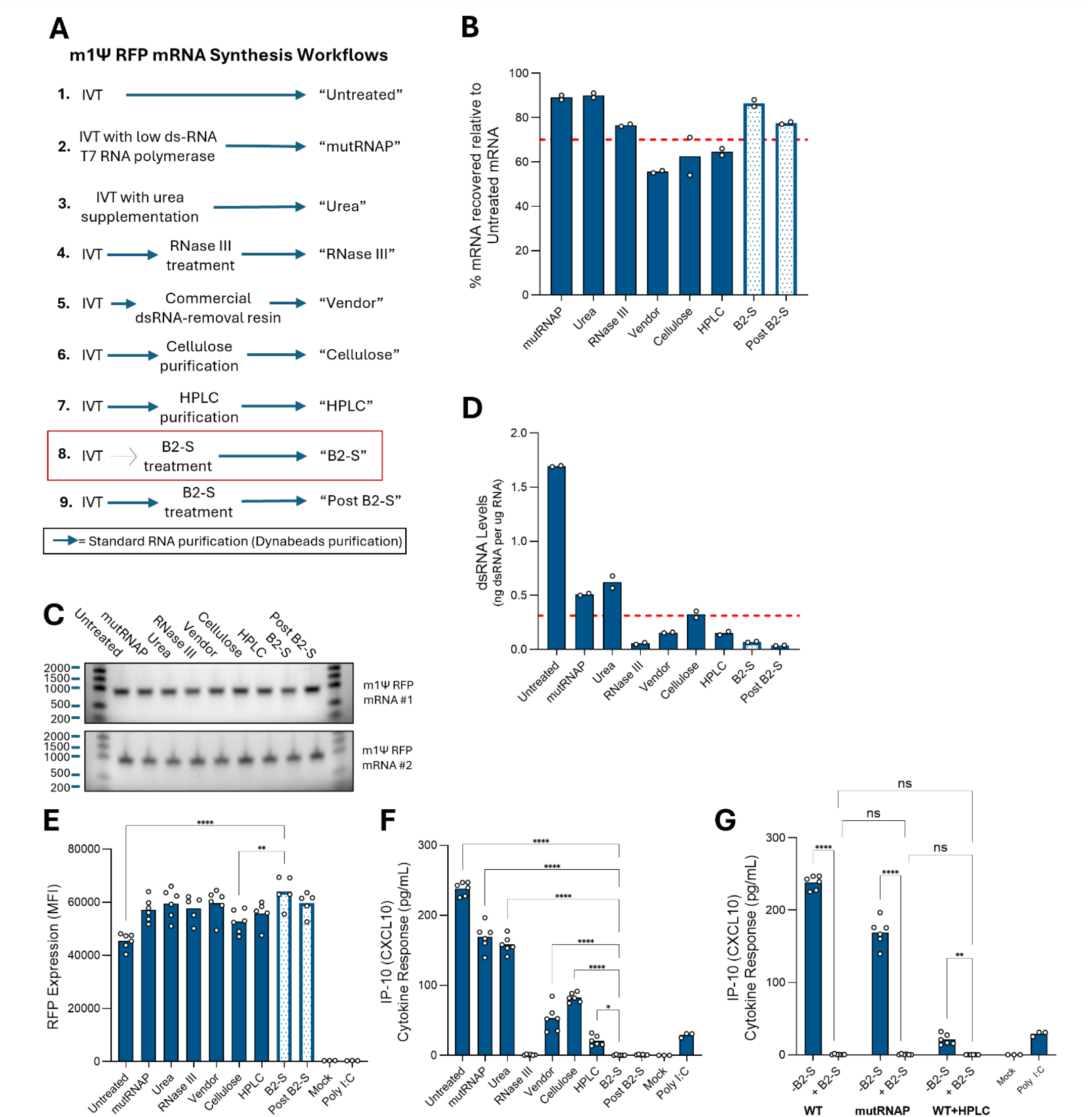
B2-S compares favorably with tested dsRNA-reduction workflows. **(A)** Schematic outlining the individual workflows for dsRNA removal and their identification names for the following plots. Each blue arrow represents a standard mRNA purification (Dynabead RNA purification; see Materials and Methods). The dotted arrow in workflow #8 indicates that the B2-S treatment began immediately on the crude IVT reaction prior to any purification. The red box surrounding workflow #8 represents our suggested workflow for dsRNA removal based on this work. **(B-D)** RFP mRNA recoveries (B), integrity (C), and dsRNA levels as measured by K1-9D5 ELISA (D) for the different workflows. The red dotted lines in (B) and (D) represent 70% recovery and the ELISA LLOQ, respectively. The numbers next to the RNA ladder in (C) represent size in nucleotides. **(E and F)** RFP expression (E) and IP- 10/CXCL10 levels in media (F) after BJ fibroblasts were transfected with RFP mRNA from the different workflows. **(G)** IP-10/CXCL10 levels in media after transfection with post-hoc B2-S treatment of mutant RNAP- or HPLC-purified mRNAs. Bars represent the mean, and individual replicate values are shown as open circles. Statistical significance was determined in (E) and (F) by one-way ANOVA followed by Šídák’s multiple comparisons test versus the B2-S sample. Comparisons that are not significant are not indicated. Statistical significance was determined in (G) by two-way ANOVA followed by Tukey’s multiple comparisons test. **\***: P≤0.05; **\*\***: P≤0.01; **\*\*\*\***: P≤0.0001, **ns**: not significant.

Relative mRNA recovery was at or above 70% for mutant RNAP, urea, RNase III, and both B2- S configurations. Cellulose, Vendor resin, and HPLC produced lower recovery (Figure 5B). None of the methods measurably affected mRNA integrity (Figure 5C). By K1-9D5 dsRNA ELISA, RNase III, Vendor resin, HPLC, and both B2-S configurations reduced dsRNA below the working threshold target (Figure 5D). The J2 dot blot was broadly consistent, although the Vendor resin sample retained detectable signal, suggesting that the ELISA and dot blot may detect different dsRNA subpopulations (Supplemental Figure 7A).

Next, we transfected mRNA from each condition into BJ fibroblasts and measured RFP expression and IP-10/CXCL10 secretion after 24 hours. The percentage of RFP-positive cells was similar across conditions (Supplemental Figure 7B). Both B2-S configurations increased RFP expression per cell relative to the untreated and cellulose conditions (Figure 5E). The most striking differences were in IP-10/CXCL10 secretion: only RNase III-treated mRNA and both B2-S-treated mRNAs reduced IP-10/CXCL10 to mock-transfection levels. Mutant RNAP, urea, cellulose, Vendor resin, and HPLC samples remained elevated, including HPLC despite passing the dsRNA ELISA target (Figure 5F). A similar but more pronounced pattern was observed in the JAWSII cell line (Supplemental Figures 7C-E). These data demonstrate that RNase III or B2-S treatment reduces dsRNA below the level required to activate cellular dsRNA sensors, and that dsRNA ELISA is necessary but not sufficient for assessing residual immunostimulatory RNA. A cell-based readout such as IP-10/CXCL10 is needed to determine whether residual RNA impurities remain biologically active.

The comparable performance of RNase III-treated and B2-S-treated RFP mRNA was unexpected, given that RNase III can exhibit activity toward unintended ssRNA substrates, even after reaction optimization (39,40). We therefore asked whether RNase III compatibility was substrate dependent. When sgRNA or EPO mRNA was subjected to the same RNase III treatment, they showed evidence of degradation, whereas B2-S treatment preserved RNA integrity (Supplemental Figure 8). These results suggest that the RFP mRNA used in the benchmark may lack RNase III-sensitive structural features under the conditions tested. More broadly, these results indicate that RNase III treatment may not be a universal solution for dsRNA depletion, because its compatibility depends on the sequence and structure of the target RNA. In contrast, B2-S achieved functional dsRNA depletion while preserving RNA integrity across multiple substrates (see Supplemental Figure 5).

Finally, we tested whether post-hoc B2-S treatment could suppress the IP-10/CXCL10 response from mutant RNAP- or HPLC-purified mRNA. B2-S treatment of either material reduced IP-10/CXCL10 to mock-transfection levels (Figure 5G and Supplemental Figures 7F- J). These results indicate that B2-S removes a biologically active dsRNA fraction that can persist after other dsRNA-control workflows, even when analytical dsRNA measurements are low. As such, B2-S can be a useful polishing step in workflows that already include other dsRNA mitigation strategies such as low dsRNA RNA polymerases.

### B2-S-treated mRNA matches HPLC-purified mRNA *in vivo* with higher recovery

We next asked how B2-S-treated mRNA would perform *in vivo*. Firefly luciferase (FLuc) m1Ψ- modified mRNA was synthesized and either B2-S-treated or HPLC-purified. mRNA recovery was almost 30% higher for B2-S-treated mRNA than for HPLC-purified mRNA with no loss of integrity (Supplemental Figures 9A and 9B). Both B2-S and HPLC reduced dsRNA well below the working threshold (Supplemental Figure 9C). In A549 cells, both conditions produced higher FLuc activity and lower IP-10/CXCL10 than untreated mRNA (Supplemental Figures 9D and 9E).

The mRNAs were then formulated into liver-targeting lipid nanoparticles (LNPs). All mRNA conditions generated homogeneous particles with high payload encapsulation (Supplemental Figures 10A and 10B). FLuc mRNA-LNPs were injected intravenously into mice, and luciferase flux was measured *ex vivo* at 4 and 24 hours to compare expression magnitude and duration (Figures 6A-C). At 4 hours, B2-S and HPLC both produced significantly higher FLuc activity than untreated mRNA, with B2-S showing the highest mean signal. At 24 hours, signal decreased in all groups, but B2-S and HPLC remained higher than untreated mRNA.

**Figure 6.**
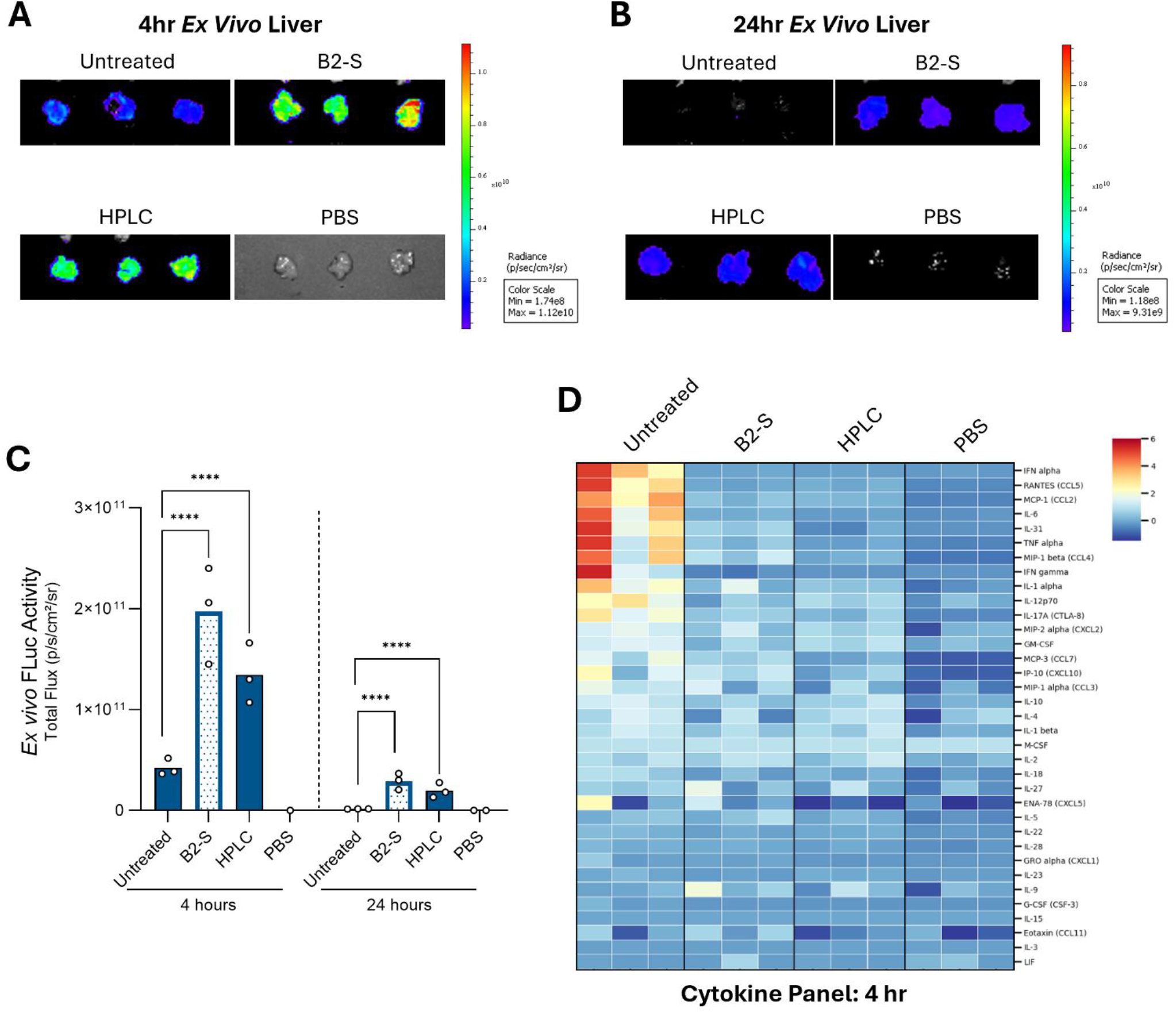
B2-S-treated mRNA matches HPLC-purified mRNA *in vivo* while suppressing inflammatory cytokine responses. (A and. **B)** Representative *ex vivo* bioluminescence images following intravenous administration of liver-targeting LNPs containing untreated, B2-S-treated, or HPLC-purified FLuc m1Ψ-modified mRNA, with PBS as a control. Luciferase activity was measured at 4 h (A) and 24 h (B) after dosing. Radiance is shown as photons/s/cm²/sr. **(C)** Quantification of *ex vivo* FLuc signal at 4 and 24 h. Bars represent the mean, and individual replicate values are shown as open circles. Statistical analysis was performed on log-transformed total flux values as described in Materials and Methods. Comparisons that are not significant are not indicated. ****: P≤0.0001. **(D)** Serum cytokine and chemokine profiles measured 4 h after dosing using a 36-plex ProcartaPlex assay. Columns represent individual animals and rows represent the indicated cytokines and chemokines; color indicates the Z-score calculated from cytokine/chemokine concentrations measured in pg/mL. N=3 per condition.

We assessed early *in vivo* toxicity by measuring levels of liver enzymes and cytokines in the serum. AST and ALT remained below 3x the upper limit of normal (ULN) in all groups, indicating no early hepatocellular injury (Supplemental Figures 10D and 10E) (41). Serum cytokine and chemokine profiling by 36-plex ProcartaPlex showed a strong inflammatory response 4 hours after dosing with untreated mRNA (Figure 6D). B2-S-treated and HPLC-purified mRNA did not upregulate cytokines, indicating that the inflammatory response was driven by mRNA payload quality rather than the LNP. Cytokine levels returned toward baseline by 24 hours (Supplemental Figure 10C). IFN-α, IFN-γ, and IL-6 followed the same pattern: elevated in the untreated mRNA group and near PBS-control levels in the B2-S and HPLC groups (Supplemental Figures 10F-H). These data show that B2-S treatment produces mRNA with *in vivo* activity and cytokine suppression comparable to HPLC-purified mRNA, but with higher recovery and a simpler workflow.

## DISCUSSION

IVT mRNA preparations contain heterogeneous dsRNA byproducts that can impair mRNA performance and confound mRNA-based experiments (10–13,18–22). dsRNA byproduct generation is sequence dependent as well as enzyme and reaction condition dependent so a solution to reduce it for one sequence may not apply to all (12–14,20,42). We show that these contaminants are difficult to quantify, biologically active at low levels, and not fully addressed by several dsRNA-depletion workflows. We also describe B2-S, an engineered dsRNA-binding protein that selectively removes dsRNA from IVT mRNA while preserving RNA integrity and recovery. Together, these results support a quality-control framework in which dsRNA burden is assessed by both analytical measurement and functional cellular response.

Accurate dsRNA quantification remains a major challenge for IVT mRNA quality assessment. J2-based dot blotting is useful for relative comparison but does not provide reliable absolute quantification. ELISA formats improve quantification, but our data show that antibody pairing and nucleotide chemistry strongly affect assay performance. J2-K2 and K1-K2 ELISAs under-detected Ψ- and m1Ψ-modified dsRNA, likely due to weak recognition by the K2 detection antibody. The K1-9D5 ELISA improved detection of modified dsRNA and provided a lower limit of quantification of 0.3125 ng dsRNA per µg RNA under the tested conditions. However, even this improved assay remained chemistry dependent, indicating that dsRNA standards should match the nucleotide composition of the mRNA sample for accurate quantification.

This is consistent with two previous reports that suggest nucleotide modification can affect these assays (25,26). The biological effects of dsRNA were also chemistry dependent. Defined dsRNA spike-in experiments showed that low dsRNA burdens reduced viability, induced TNFα secretion, and suppressed GFP expression. The relative disruptive potency followed the trend Ψ > U > m1Ψ > 5moU in the tested JAWSII model. To our knowledge, the cellular effects of both the amount and nucleotide chemistry of defined dsRNA impurities in a mRNA preparation have not been systematically characterized. The finding that Ψ-dsRNA was more disruptive than unmodified dsRNA was unexpected because Ψ modification usually reduces innate immune activation in IVT mRNA (23). This difference may reflect how modified nucleotides alter dsRNA structure, thereby changing stability, nuclease sensitivity, or helix geometry, which in turn, may affect dsRNA-sensor activation. Further studies will be needed to define the mechanism.

The comparison between m1Ψ- and 5moU-modified mRNA further shows that reduced immunostimulation does not predict higher mRNA activity. In the spike-in assay, 5moU- modified dsRNA was the least inflammatory species. However, after HPLC purification reduced dsRNA below the working analytical target, m1Ψ-modified mRNA produced higher protein expression than 5moU-modified mRNA without a marked increase in TNFα secretion. This suggests that 5moU may reduce cytokine-inducing RNA-sensor activation but limit translational performance. Overall, the data support m1Ψ modification combined with effective dsRNA removal as a strong strategy for achieving high mRNA expression with low innate immune activation, consistent with previous work showing improved function of m1Ψ-modified mRNA following dsRNA purification (11,19,35).

Our work shows that dsRNA-reduction workflows are not equivalent and that each approach involves distinct trade-offs. HPLC is widely regarded as the benchmark method for dsRNA removal, but it does not specifically recognize dsRNA (10,17). Rather, separation is achieved through physicochemical differences, which can limit selectivity and contribute to reduced recovery. Likewise, approaches that decrease dsRNA formation during IVT, including engineered RNA polymerases (13,20) or reaction optimization (21,22), can substantially lower dsRNA burden but are unlikely to eliminate it completely. Enzymatic approaches such as RNase III can be highly effective (19) but can also introduce risks to RNA integrity. Consistent with our observation that RNase III degraded sgRNA and EPO mRNA, recent work showed that RNase III can extensively degrade single-stranded mRNA, even after reaction optimization (39,40). Consistent with these limitations, several workflows in our benchmark study lowered dsRNA signal by ELISA, but not all suppressed the IP-10/CXCL10 cytokine response after transfection. HPLC-purified and RNase III-treated mRNA both met the working analytical threshold, yet only the HPLC-purified mRNA retained measurable IP- 10/CXCL10 induction. This disconnect suggests that ELISA-recognized dsRNA burden does not always predict residual immunostimulatory activity. Therefore, IP-10/CXCL10 serves as a useful orthogonal readout because it reflects activation of downstream antiviral interferon signaling triggered by dsRNA sensing, rather than antibody recognition of a specific RNA species (37). Together, these findings highlight the value of pairing analytical dsRNA quantification with a functional cell-based assay.

The choice of functional assay is important. Engineered reporter cell lines, such as A549-Dual (InvivoGen), help provide convenient readouts of defined innate immune signaling events, such as interferon regulatory factor activation, but their response depends on the signaling pathways present in the cell line and the specific reporter being measured, which may not fully reflect endogenous responses to residual immunostimulatory RNA. In this study, BJ fibroblasts and JAWSII cells provided complementary dsRNA-responsive models in which endogenous cytokine production could be measured directly. BJ fibroblasts were particularly useful for monitoring IP-10/CXCL10 as a downstream interferon-responsive marker (35,37), whereas JAWSII cells provided a sensitive innate-immune background for assessing cytokine induction, viability, and mRNA expression (36,43). Together, these models allowed residual immunostimulatory activity to be assessed across distinct cellular contexts rather than through a single engineered reporter pathway readout.

Our B2-S dsRNA removal system addresses these limitations through a simple affinity-based workflow and compares favorably against current dsRNA-reduction strategies. The engineered single-chain design of the protein preserves dsRNA binding while reducing the complexity of the dimeric scaffold. B2-S displays several properties that are well suited for selective dsRNA capture: it binds duplexes of approximately 21 bp or longer, is sensitive to internal base-pair mismatches, and shows minimal detectable binding to structured ssRNA. These features may help limit recognition of mRNA secondary structures, which frequently contain bulges, mismatches, and relatively short duplex regions (33). In our benchmark experiments, B2-S reduced biologically active dsRNA while preserving mRNA recovery, integrity, and function. Compared with HPLC, B2-S-treated mRNA showed comparable functional performance with higher mRNA recovery. Together, these results support B2-S as a robust, non-chromatographic, accessible, and yield-preserving dsRNA-removal strategy that produces highly active mRNA with low immunostimulatory activity. Across the samples tested, B2-S treatment consistently reduced IP-10/CXCL10 induction to levels comparable to transfection controls. Importantly, B2-S-treated samples met both the analytical dsRNA target and the functional IP-10/CXCL10 benchmark, a relationship that was not observed for several other dsRNA-reduction workflows. Only RNase III showed similar performance in the benchmark comparison; however, it introduces risks to RNA integrity (39,40). With further validation across a broader range of transcripts and process conditions, this relationship could support using both the dsRNA ELISA and cell-based cytokine assays to qualify an IVT workflow with B2-S treatment. Once the process is established, analytical dsRNA measurement alone may be sufficient for routine assessment of B2-S-treated samples.

Our results show that residual dsRNA can also affect repeat mRNA delivery. Untreated EPO mRNA produced early protein expression but failed to sustain expression after repeat transfection and induced IP-10/CXCL10. B2-S-treated EPO mRNA maintained expression after the second dose while preserving viability and normal cell morphology. These findings have implications beyond mRNA therapeutics. IVT mRNA is widely used as a research reagent for transient protein expression, genome editing, delivery optimization, reprogramming, and cell-state manipulation (1,5–7). In these settings, residual dsRNA can act as an experimental confounder. Changes in expression, editing efficiency, cytokine secretion, viability, morphology, or transcriptional state may reflect dsRNA-driven innate immune activation rather than the intended effect of the mRNA payload or delivery condition.

In summary, dsRNA contaminants in IVT mRNA are analytically challenging, biologically active, and capable of confounding mRNA function. Modified dsRNA species can be under-detected by common antibody-based assays, and analytical dsRNA depletion does not always predict suppression of cellular innate immune activation. B2-S provides a selective, non-chromatographic, yield-preserving workflow that removes biologically active dsRNA while maintaining mRNA integrity. These findings support paired analytical and cell-based assays for IVT mRNA quality assessment and establish B2-S as an accessible and fast approach for producing mRNA with improved expression and reduced immunostimulatory activity.

## ACKNOWLEDGEMENTS

We thank Meg Rogge for reviewing the manuscript. We also thank Kyle Morris, Xiaohua Xu, and Xiquan Liang for their assistance with *E. coli* fermentation and protein purification.

## AUTHOR CONTRIBUTIONS

Tyson Vonderfecht: *Conceptualization, Formal Analysis, Investigation, Methodology, Project Administration, Resources, Validation, Visualization, Writing—original draft & review editing.* Peter Lam: *Formal Analysis, Investigation, Resources, Visualization, Writing—review & editing.* Genia Verovskaya: *Formal Analysis, Investigation, Methodology, Supervision, Writing—review & editing.* Jason Potter: *Conceptualization, Funding Acquisition, Supervision, Writing—review & editing*.

## SUPPLEMENTARY DATA

Supplementary Data are available online.

## DATA AVAILABILITY

The data underlying this article will be shared on reasonable request to the corresponding authors and may be subject to applicable material-transfer and intellectual-property restrictions. The B2-S construct is subject to intellectual property restrictions.

## SUPPLEMENTAL INFORMATION

**Supplemental Figure 1.**
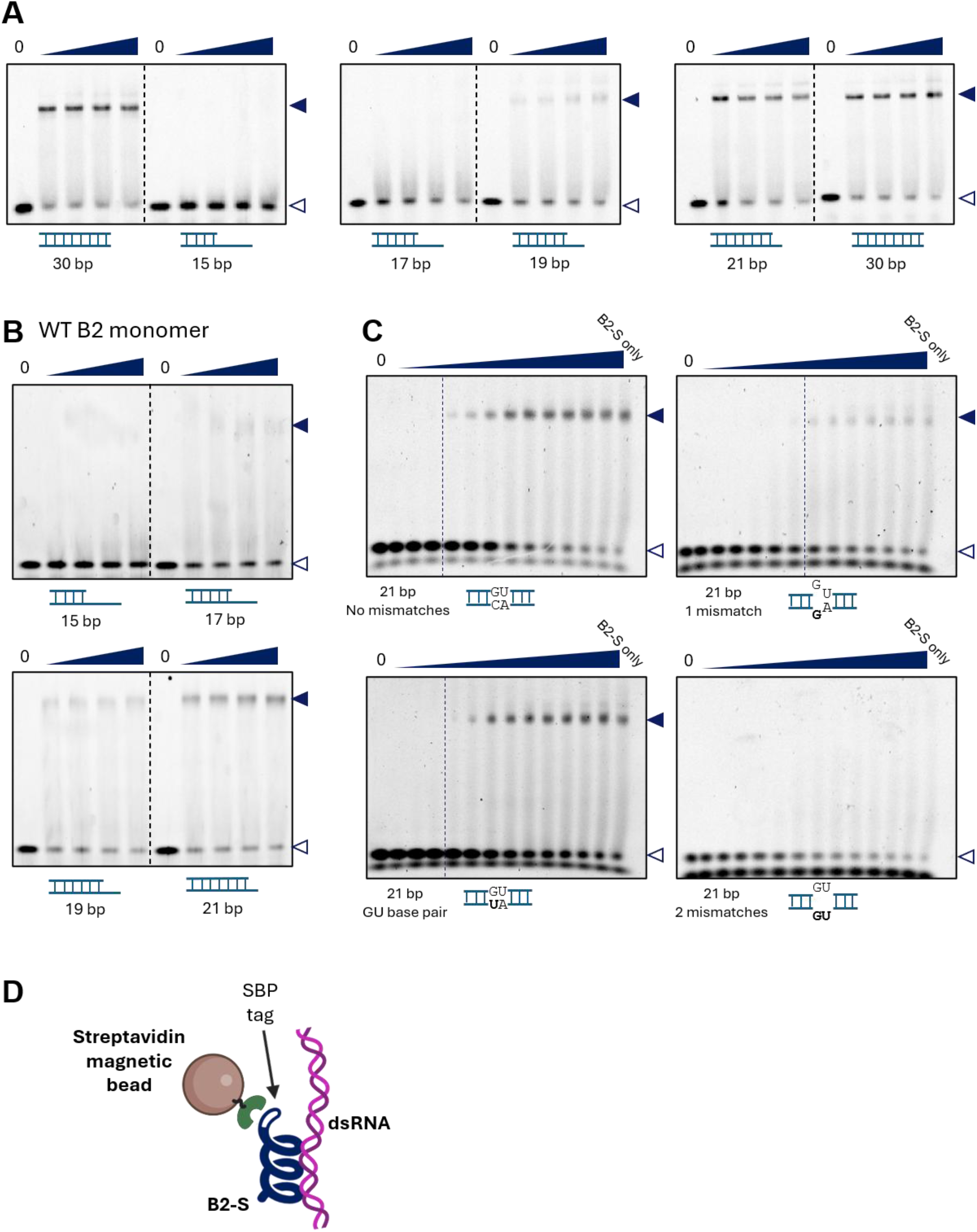
B2-S preferentially binds perfectly matched dsRNA substrates that are ≥21 bp. **(A and B)** Electrophoretic mobility shift assays (EMSAs) of B2-S (A) or wild-type (WT) B2 monomer (B) with 15-, 17-, 19-, 21-, or 30-bp dsRNA substrates. **(C)** EMSAs of B2-S with a 21-bp dsRNA substrate containing either no mismatch, one mismatch, two mismatches, or a central GU base pair. For each assay in (A–C), a constant amount of dsRNA substrate was incubated with increasing concentrations of B2-S or B2 monomer, as indicated. “B2-S only” lanes contain protein in the absence of dsRNA, whereas “0” lanes contain dsRNA substrate in the absence of protein. Filled arrowheads indicate bound protein–dsRNA complexes, and open arrowheads indicate free dsRNA substrate. **(D)** Schematic illustrating the use of B2-S to capture and remove dsRNA. SBP, streptavidin-binding peptide. Image was made with BioRender.

**Supplemental Figure 2.**
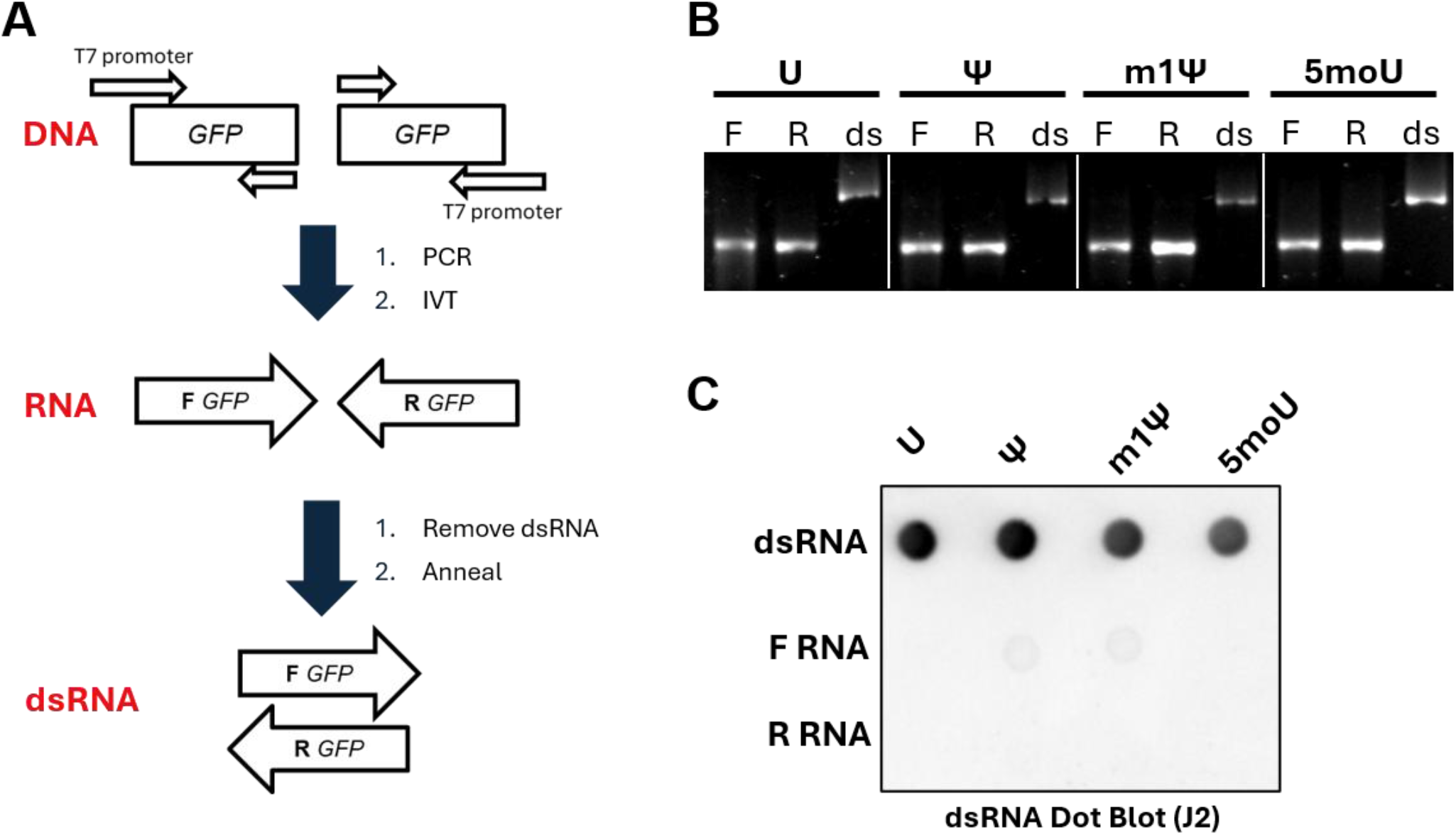
Generation and validation of dsRNA. **(A)** Schematic illustrating the workflow used to generate dsRNA. DNA templates containing a T7 promoter were amplified using forward and reverse primers and used for *in vitro* transcription (IVT) to generate complementary forward (F) and reverse (R) RNA strands. Residual dsRNA byproducts were removed from the individual RNA preparations by cellulose–ethanol purification, and the complementary RNAs were subsequently annealed to generate dsRNA. **(B)** Agarose gel analysis of RNA products generated using the indicated modified nucleotides: unmodified U (U), pseudouridine (Ψ), N1-methylpseudouridine (m1Ψ), or 5- methoxyuridine (5moU). F, forward RNA; R, reverse RNA; ds, annealed dsRNA. The slower-migrating band indicates formation of the dsRNA product. **(C)** Dot-blot analysis of the corresponding dsRNA preparations, confirming successful generation of dsRNA containing each indicated uridine modification.

**Supplemental Figure 3.**
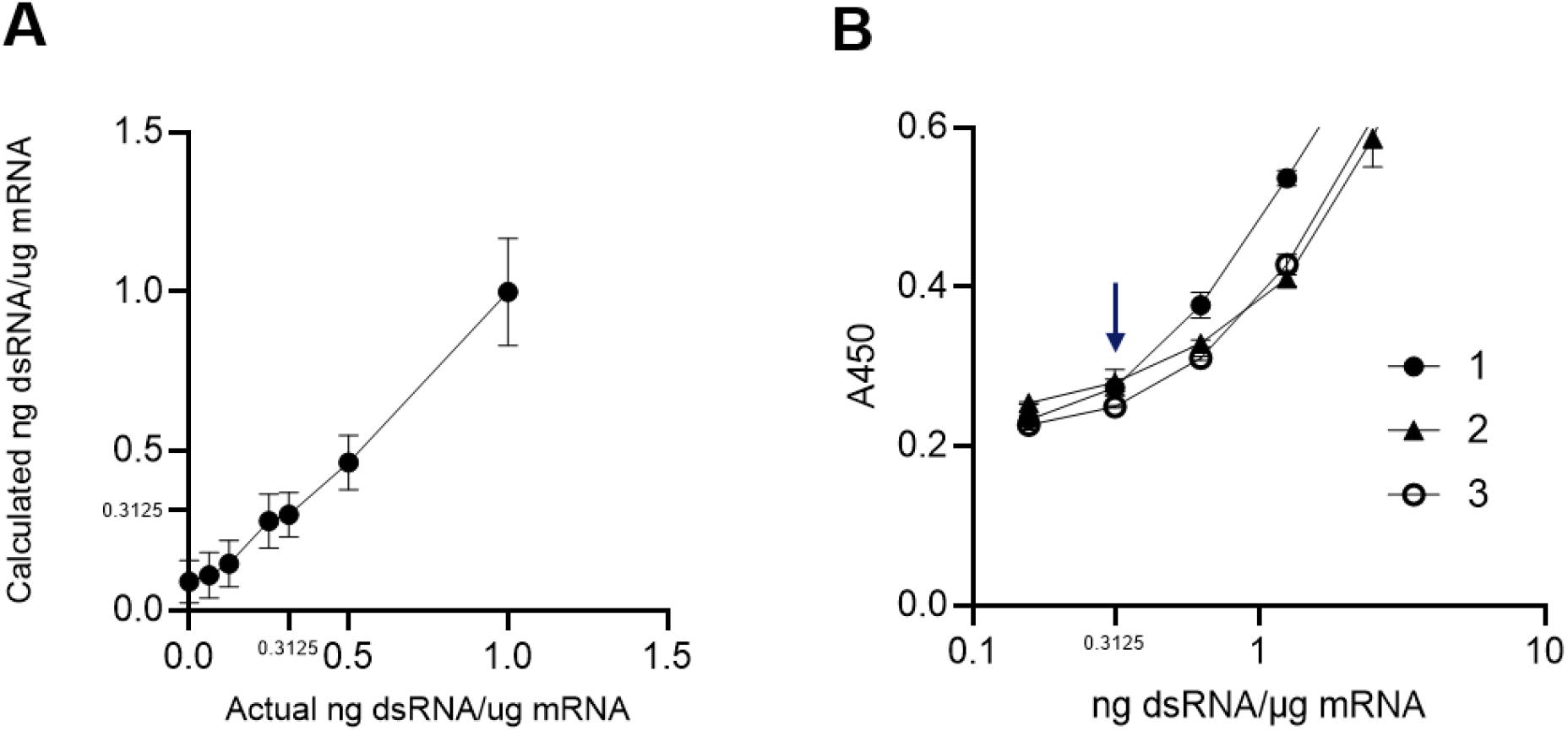
Determination of the lower limit of quantification (LLOQ) for the K1-9D5 ELISA. **(A)** HPLC-purified m1Ψ-modified RFP or Cas9 mRNA was spiked with known amounts of m1Ψ-modified dsRNA and analyzed by K1-9D5. K1-9D5 measurements closely matched expected dsRNA concentrations down to 0.3125 ng dsRNA per µg RNA, whereas quantification at lower spike-in levels was increasingly affected by assay background. **(B)** Standard curves from three independent K1-9D5 ELISA runs. The arrow indicates 0.3125 ng dsRNA per µg RNA, corresponding to the transition from the low-signal region to the responsive portion of the curve and defined as the K1-9D5 LLOQ.

**Supplemental Figure 4.**
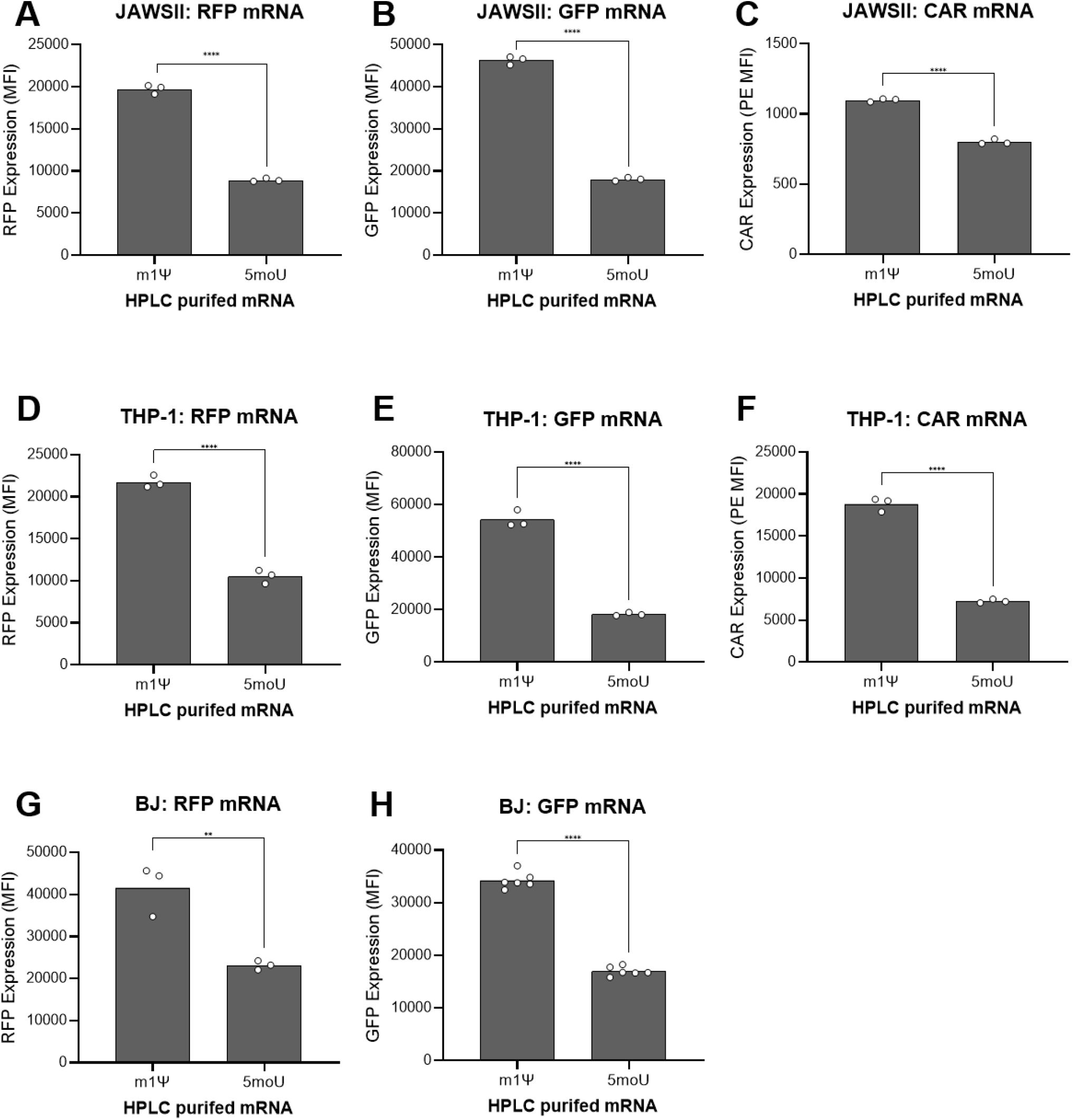
Enhanced expression of HPLC-purified m1Ψ-modified mRNA extends across multiple constructs and cell types. HPLC-purified mRNAs containing either m1Ψ or 5moU were evaluated for protein expression in multiple cellular contexts. **(A–C)** RFP, GFP, and CD19-CAR expression in murine JAWSII immature dendritic cells. **(D–F)** RFP, GFP, and CD19-CAR expression in human THP-1 monocytes. **(G–H)** RFP and GFP expression in human BJ fibroblast cells. Bars represent the mean fluorescence intensity (MFI) and individual points represent replicates. Statistical significance was determined by unpaired t-test. **\*\***: P≤0.01; **\*\*\*\***: P≤0.0001.

**Supplemental Figure 5.**
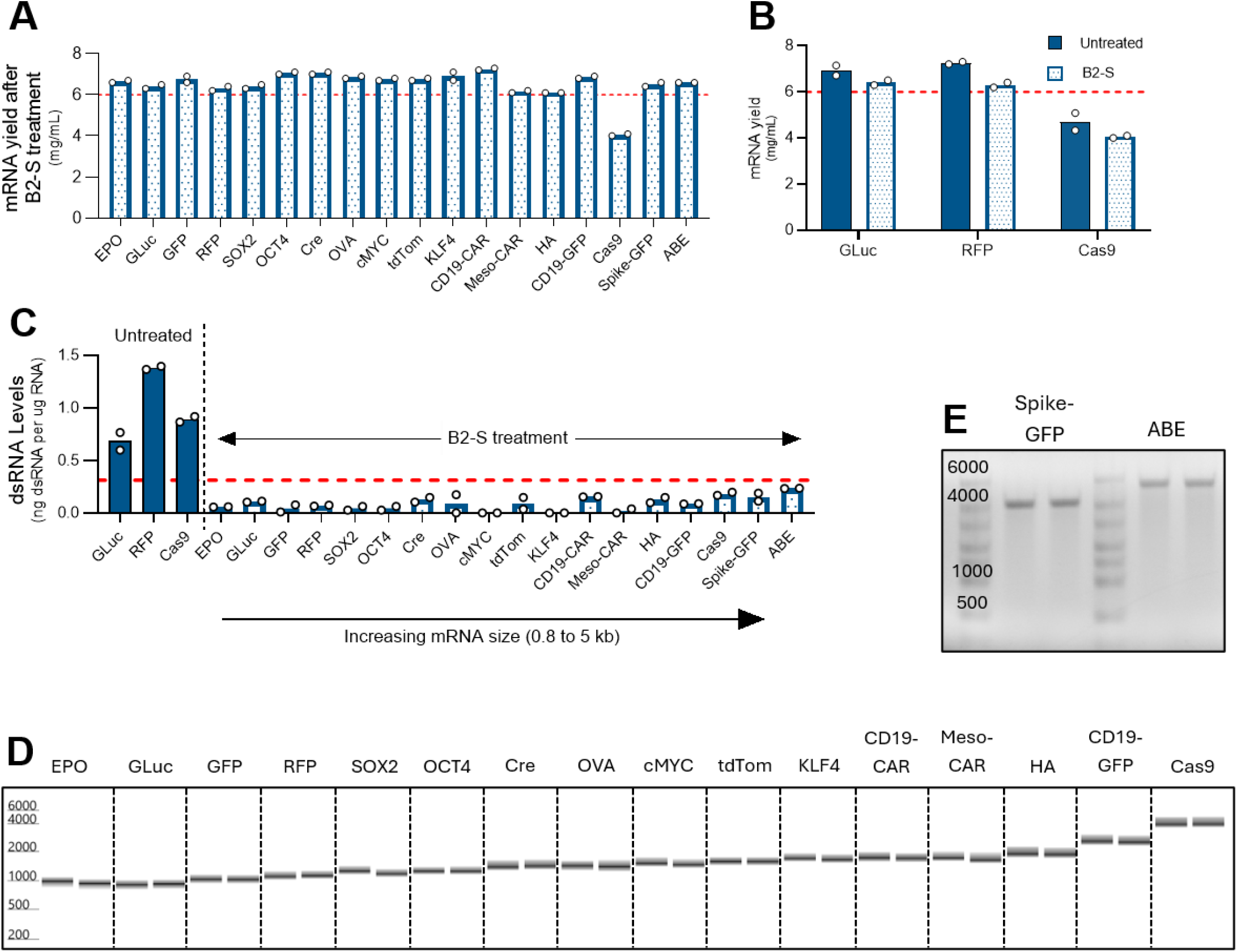
B2-S treatment efficiently removes dsRNA while preserving mRNA yield and integrity across diverse IVT mRNA constructs. **(A)** Recovery of IVT mRNA following B2-S treatment across multiple mRNA constructs. The dashed red line indicates an IVT mRNA yield of 6 mg/mL. **(B)** Comparison of mRNA recovery before and after B2-S treatment for selected constructs. The B2-S-treated Cas9 mRNA remained below 6 mg/mL because the corresponding untreated mRNA also had a starting yield below 6 mg/mL, rather than a substantial yield loss during treatment. The dashed red line indicates an IVT mRNA yield of 6 mg/mL. **(C)** dsRNA levels in untreated and B2-S-treated mRNA samples measured by the K1-9D5 ELISA. The dashed red line indicates the lower limit of quantification of 0.3125 ng dsRNA per µg mRNA. B2-S treatment reduced dsRNA to below this threshold across the tested constructs. **(D and E)** Gel-like images from capillary electrophoresis (D) or agarose gel electrophoresis (E) showing mRNA integrity across constructs ranging from approximately 0.8 to 5 kb following B2-S treatment. Bars in plots represent the mean and individual points represent replicate measurements.

**Supplemental Figure 6.**
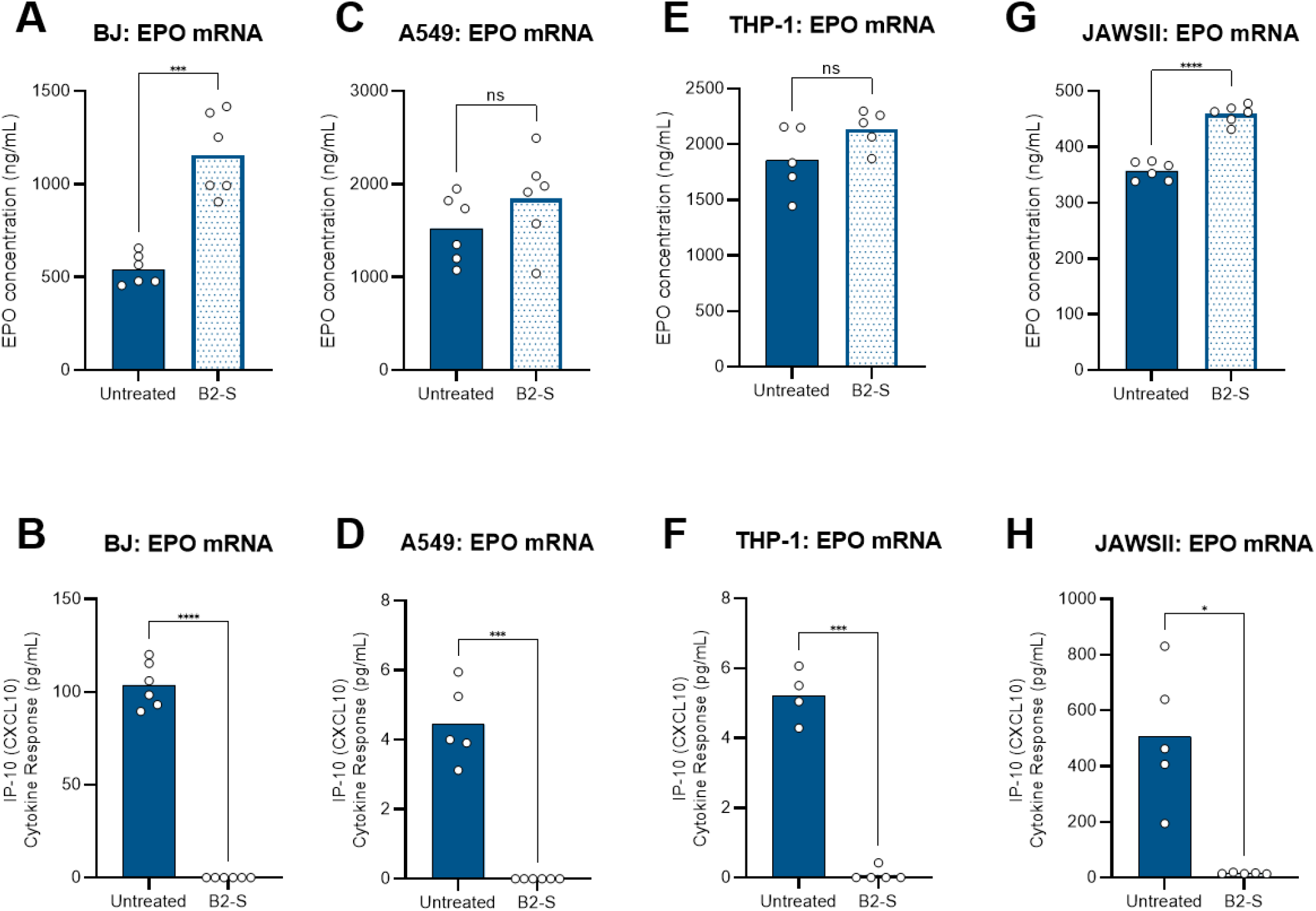
B2-S treatment improves EPO expression while reducing IP- 10/CXCL10 induction across multiple cell lines. **(Top panels: A, C, E, and G)** EPO expression following transfection with untreated or B2-S-treated EPO mRNA in the indicated cell lines. **(Bottom panels: B, D, F, and H)** IP-10/CXCL10 levels measured in the corresponding samples. B2-S treatment increased EPO expression for most cell lines while markedly reducing IP-10/CXCL10 induction across all the tested cell lines. Bars represent the mean, and individual points represent replicate measurements. Statistical significance was determined by unpaired t-test. **\***: P≤0.05; **\*\*\***: P≤0.001; **\*\*\*\***: P≤0.0001, **ns**: not significant.

**Supplemental Figure 7.**
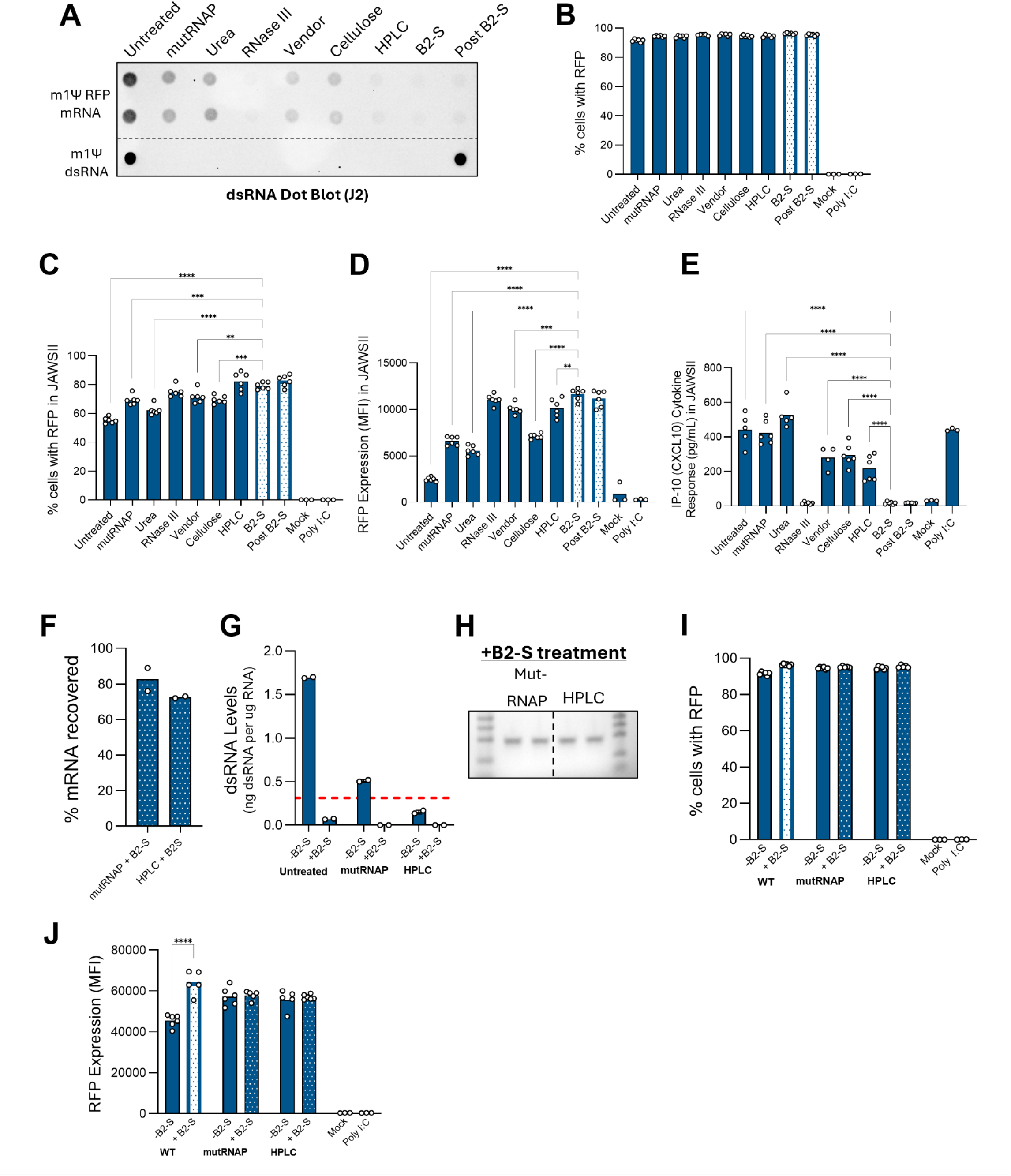
Functional and analytical comparison of dsRNA-reduction workflows and post-hoc B2-S treatment. **(A)** J2 dot blot analysis of residual dsRNA in RFP mRNA prepared using the indicated dsRNA-reduction workflows. **(B)** Percentage of RFP-positive BJ fibroblasts 24 h after transfection with RFP mRNA generated using the indicated workflows. **(C–E)** Functional evaluation of the same RFP mRNA preparations in JAWSII cells, including the percentage of RFP-positive cells (C), RFP expression (D), and IP-10/CXCL10 secretion (E). **(F–J)** Characterization of mutant RNAP- and HPLC-purified RFP mRNA before and after post-hoc B2-S treatment. mRNA recovery (F), residual dsRNA measured by K1-9D5 ELISA (G), mRNA integrity (H), percentage of RFP-positive cells (I), and RFP expression (J) were evaluated in BJ fibroblasts following B2-S treatment. The dashed red line in (G) indicates the K1-9D5 lower limit of quantification (LLOQ) of 0.3125 ng dsRNA per µg mRNA. Mock-transfected and poly(I:C)-treated cells were included as negative and positive controls, respectively, where indicated. Bars represent the mean, and individual replicate values are shown as open circles. Statistical significance was determined in (C-E) by one-way ANOVA followed by Dunnett’s multiple comparisons test versus the B2-S sample and in (I-J) by two-way ANOVA followed by Tukey’s multiple comparisons test versus the respective B2-S-treated sample. Not significant comparisons are not indicated. **\*\***: P≤0.01; **\*\*\***: P≤0.001; **\*\*\*\***: P≤0.0001.

**Supplemental Figure 8.**
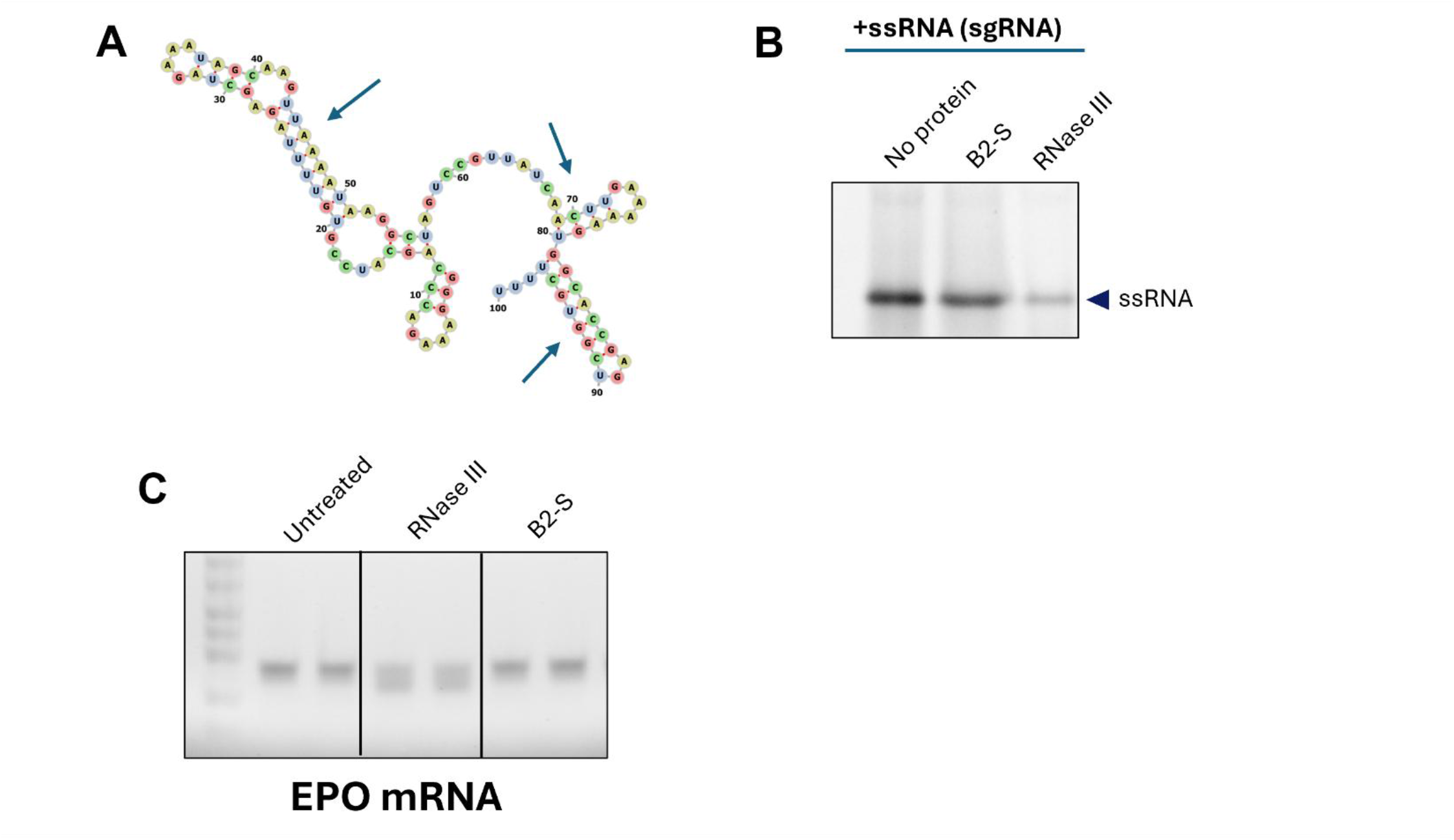
RNase III sensitivity is substrate dependent, whereas B2-S treatment preserves RNA integrity. **(A)** Predicted secondary structure of the sgRNA used in this study. See Supplemental Methods for the sequence. Nucleotides are color coded by base identity: adenine (A), yellow; uracil (U), blue; guanine (G), red; and cytosine (C), green. Arrows indicate intramolecular base-paired regions with dsRNA-like structure. Image was generated with RNAfold (44). **(B)** Gel analysis of sgRNA following the indicated treatments, showing degradation after RNase III treatment compared with preservation of the intact RNA species following B2-S treatment. **(C)** Gel analysis of EPO mRNA following the indicated treatments, demonstrating RNase III-associated degradation and preservation of mRNA integrity following B2-S treatment.

**Supplemental Figure 9.**
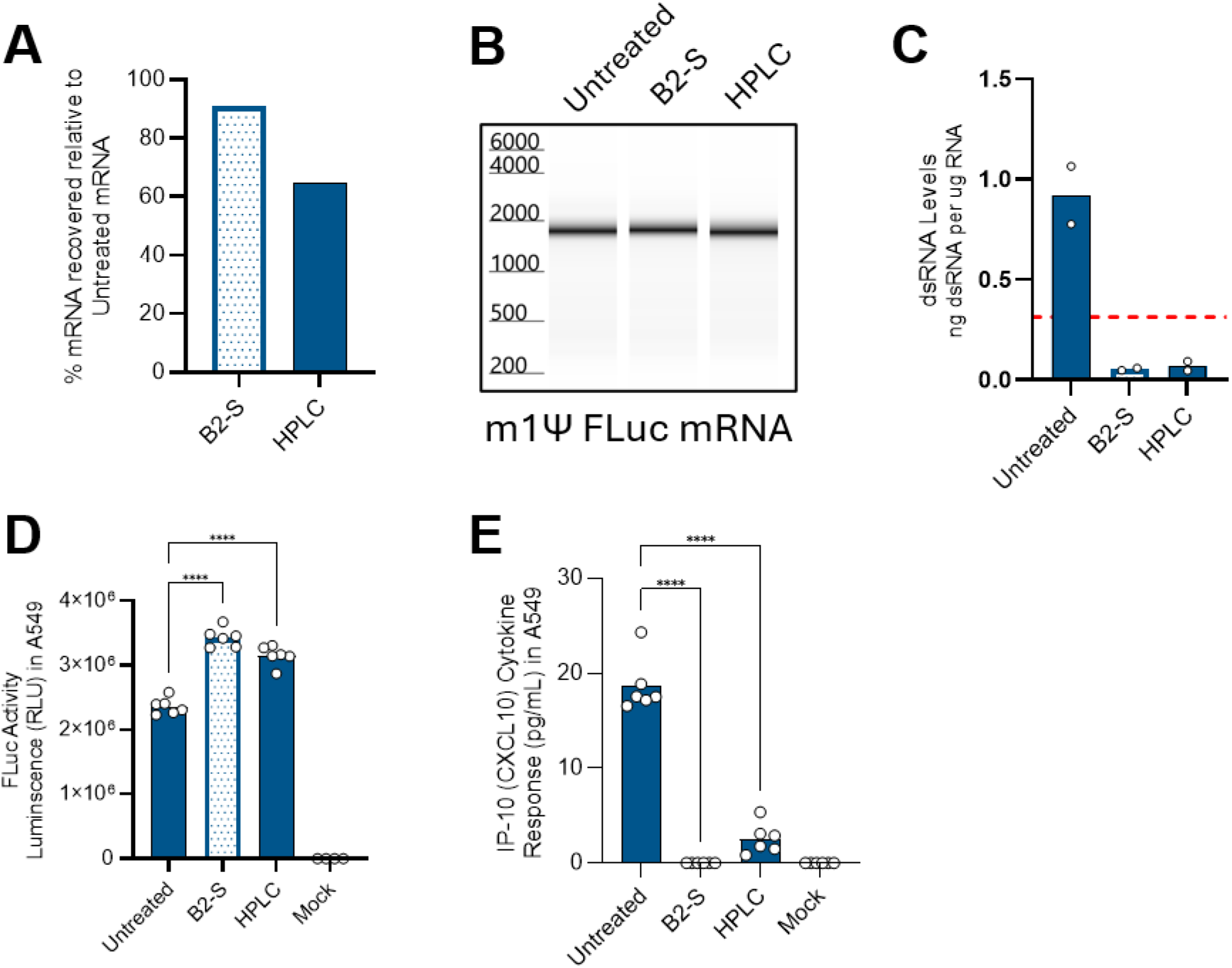
B2-S treatment preserves FLuc mRNA recovery and integrity while reducing dsRNA and improving functional performance. **(A)** Recovery of m1Ψ- modified firefly luciferase (FLuc) mRNA following B2-S treatment or HPLC purification relative to untreated mRNA. **(B)** Capillary electrophoresis analysis showing preservation of FLuc mRNA integrity following B2-S treatment and HPLC purification. **(C)** Residual dsRNA levels in untreated, B2-S-treated, and HPLC-purified FLuc mRNA measured by the K1-9D5 ELISA. The dashed red line indicates the assay LLOQ. **(D)** FLuc activity in A549 cells following transfection with untreated, B2-S-treated, or HPLC-purified FLuc mRNA. **(E)** IP-10/CXCL10 levels in culture media from the corresponding A549 cell samples. Bars represent the mean, and individual replicate values are shown as open circles. Statistical significance was determined by one-way ANOVA followed by Tukey’s multiple comparisons test versus the B2-S sample. Not significant comparisons are not indicated. **\*\*\*\***: P≤0.0001.

**Supplemental Figure 10.**
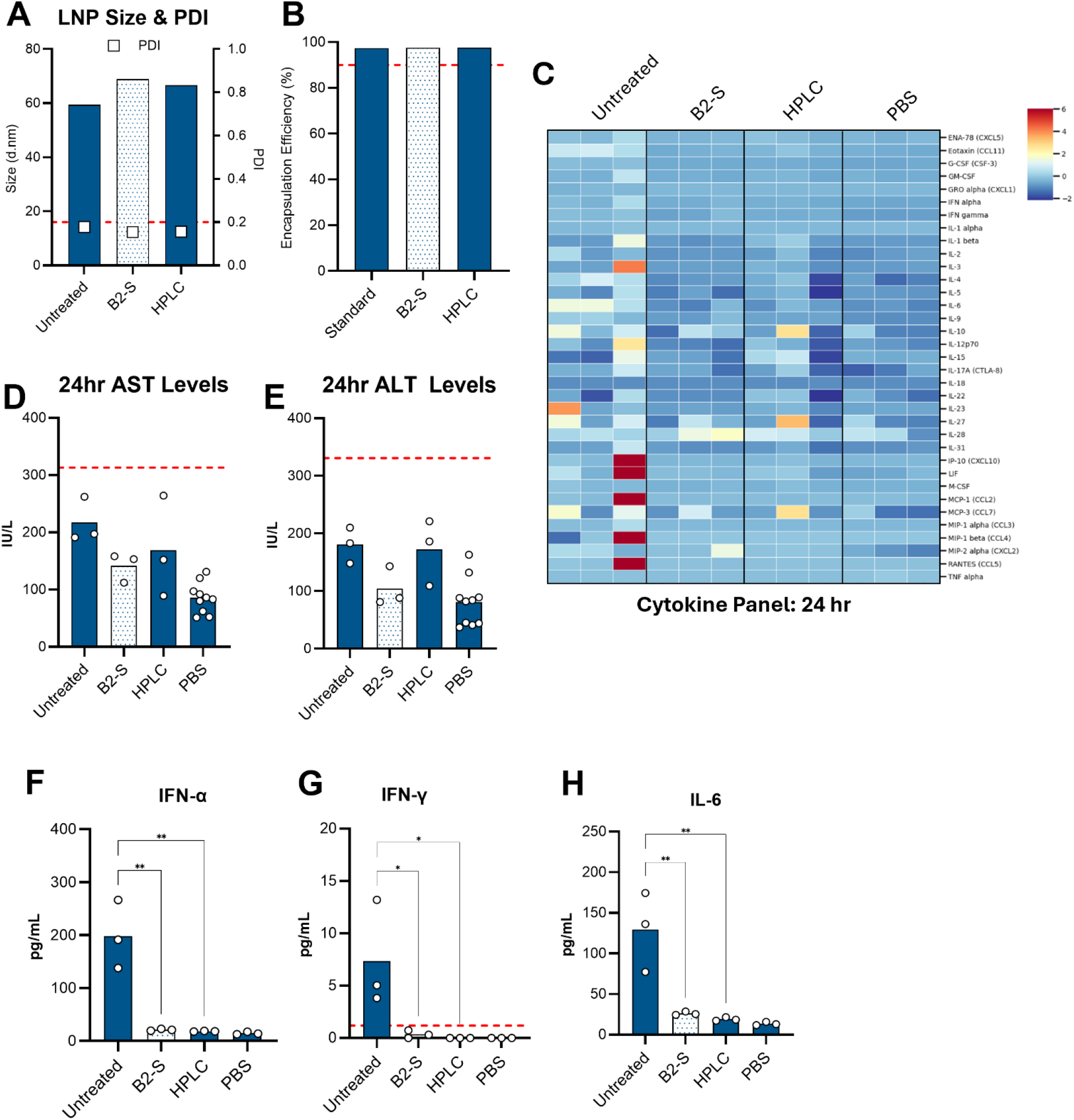
B2-S-treated mRNA shows low *in vivo* toxicity and inflammatory signaling. **(A)** Particle size and polydispersity index (PDI) of LNPs formulated with untreated, B2-S-treated, or HPLC-purified FLuc mRNA. Particle size is plotted on the left y-axis and PDI on the right y-axis. **(B)** Encapsulation efficiency of FLuc mRNA in the corresponding LNP formulations. The red dotted line indicates 90%. **(C)** Serum cytokine and chemokine profiles measured by 36-plex ProcartaPlex at 24 h after intravenous administration of FLuc mRNA-LNPs. Values are displayed as Z-scores (pg/mL) for each analyte across treatment groups. **(D and E)** Serum aspartate aminotransferase (AST) (D) and alanine aminotransferase (ALT) (E) levels following LNP administration. Dashed red lines indicate three times the upper limit of normal (3× ULN). **(F–H)** Serum IFN-α (F), IFN-γ (G), and IL-6 (H) levels 4 hours following administration of untreated, B2-S-treated, or HPLC-purified FLuc mRNA-LNPs. The dashed red line in (G) indicates the lower limit of detection (LLOD) for IFN-γ of 1.2 pg/mL. PBS-treated animals are included as controls where indicated. Bars represent the mean, and individual animals are shown as open circles. Statistical significance was determined by one-way ANOVA followed by Tukey’s multiple comparisons test. Not significant comparisons are not indicated. **\***: P≤0.05; **\*\***: P≤0.01.

## SUPPLEMENTAL METHODS

### dsRNA substrate preparation and B2 binding assays

Short dsRNA substrates were generated by annealing complementary synthetic RNA oligonucleotides. For the dsRNA length-series experiments, a common forward RNA oligonucleotide (5’-CUGACACAACUGUGUUCACUAGCAACCUCA-3’) containing a 3’ Alexa Fluor 488 label was annealed to complementary reverse oligonucleotides to generate 15- (5′- ACACAGUUGUGUCAG-3′), 17- (5’-GAACACAGUUGUGUCAG-3’), 19- (5′- GUGAACACAGUUGUGUCAG-3′), 21- (5′-UAGUGAACACAGUUGUGUCAG-3’), or 30-bp (5′- UGAGGUUGCUAGUGAACACAGUUGUGUCAG-3′) duplexes.

For analysis of the effects of mismatches and G-U base pairing, 21-bp unlabeled RNA substrates were prepared using the following oligonucleotides: 21-F, 5’- GCGUACGUACGUACGUACGUG-3’; 21-R, 5’-CACGUACGUACGUACGUACGC-3’; 21-R1, 5’- CACGUACGUAGGUACGUACGC-3’; 21-R2, 5’-CACGUACGUUGGUACGUACGC-3’; and 21-R1-U, 5’-CACGUACGUAUGUACGUACGC-3’. The corresponding forward and reverse oligonucleotides were combined to generate a perfectly matched 21-bp duplex (21-F + 21- R), substrates containing one (21-F + 21-R1) or two (21-F + 21-R2) internal mismatches, or a substrate containing an internal G-U base pair (21-F + 21-R1-U), as indicated. All RNA oligos were purchased from IDT.

For preparation of the unlabeled 21-bp substrates, RNA oligonucleotides were resuspended in water at 100 µM, and concentrations were measured by NanoDrop. Oligonucleotides were diluted to 50 ng/µL, and 25 µL of the forward oligonucleotide and 25 µL of the corresponding reverse oligonucleotide were combined with 12.5 µL of 5× annealing buffer. Samples were incubated at 70°C for 5 min, after which the heat block was turned off and the samples were allowed to cool gradually to room temperature. The resulting dsRNA preparations had a nominal concentration of 20 ng/µL.

For electrophoretic mobility-shift assays, each 10-µL binding reaction contained 10 ng dsRNA substrate, 1X B2-S binding buffer, and 4 µL of the indicated B2 protein dilution. B2 proteins were prepared as two-fold serial dilutions in 1X binding buffer. Protein and RNA were incubated under the binding conditions described in the main Materials and Methods, and protein–RNA complexes were subsequently analyzed by native gel electrophoresis.

### Primers for generation of the reverse-strand dsRNA template

Forward primer:

5’-TAATACGACTCACTATAAGGAGATGCCGCCCACTCAGACTTTATTCAAAGACC-3’

Reverse primer:

5’-AGGAGAACTCTTCTGGTCCCCACAG-3’

### THP-1/A549 cell culture and transfection

The human THP-1 monocytic and A549 lung carcinoma cell lines were obtained from the American Type Culture Collection (ATCC) and cultured according to ATCC recommendations. All media and supplements were obtained from Thermo Fisher.

20,000 cells were seeded in 100 µL media onto 96-well flat bottom cell culture plates (Thermo Fisher) for mRNA transfections. Cells in each well were transfected with 200 ng mRNA by Lipofectamine™ MessengerMAX™ Transfection Reagent (Thermo Fisher) by following the manufacturer’s 0.3-µL protocol.

### Cell assays and data analysis

Supplemental Figures 4C and 4F use a CAR construct with a V5 epitope tag. To measure CAR expression by flow cytometry, cells were stained with the V5 Tag Monoclonal Antibody (TCM5), PE (12-6796-42, Thermo Fisher) according to the manufacturer’s instructions. PE fluorescence was detected in the YL1 channel.

The levels of IP-10/CXCL10 in 50 µL JAWSII cell media were measured with the Mouse IP-10 (CXCL10) ELISA Kit (BMS6018, Thermo Fisher) by following the manufacturer’s instructions. Absorbances were read by the Varioskan microplate reader. All standard curves were constructed as described in Materials and Methods. FLuc activity was measured using the Invitrogen Luc-Screen™ Extended-Glow Luciferase Reporter Gene Assay System (T1033, Thermo Fisher) by following the manufacturer’s instructions. Luminescence was read by the Varioskan microplate reader.

### sgRNA sequence for RNAfold

5’-GGGAAAGACCCAGCAUCCGUGUUUUAGAGCUAGAAAUAGCAAGUUAAAAUAAGGCUAGUCCGUUAUCAACUUGAAAAAGUGGCACCGAGUCGGUGCUUUUU-3’

The underlined portion is the gRNA spacer.

