## Supplemental data and information for "Analytical and functional clearance of dsRNA contaminants from *in vitro* transcribed mRNA by an engineered dsRNA-binding protein"

### MANUSCRIPT TITLE

### SUPPLEMENTARY DATA

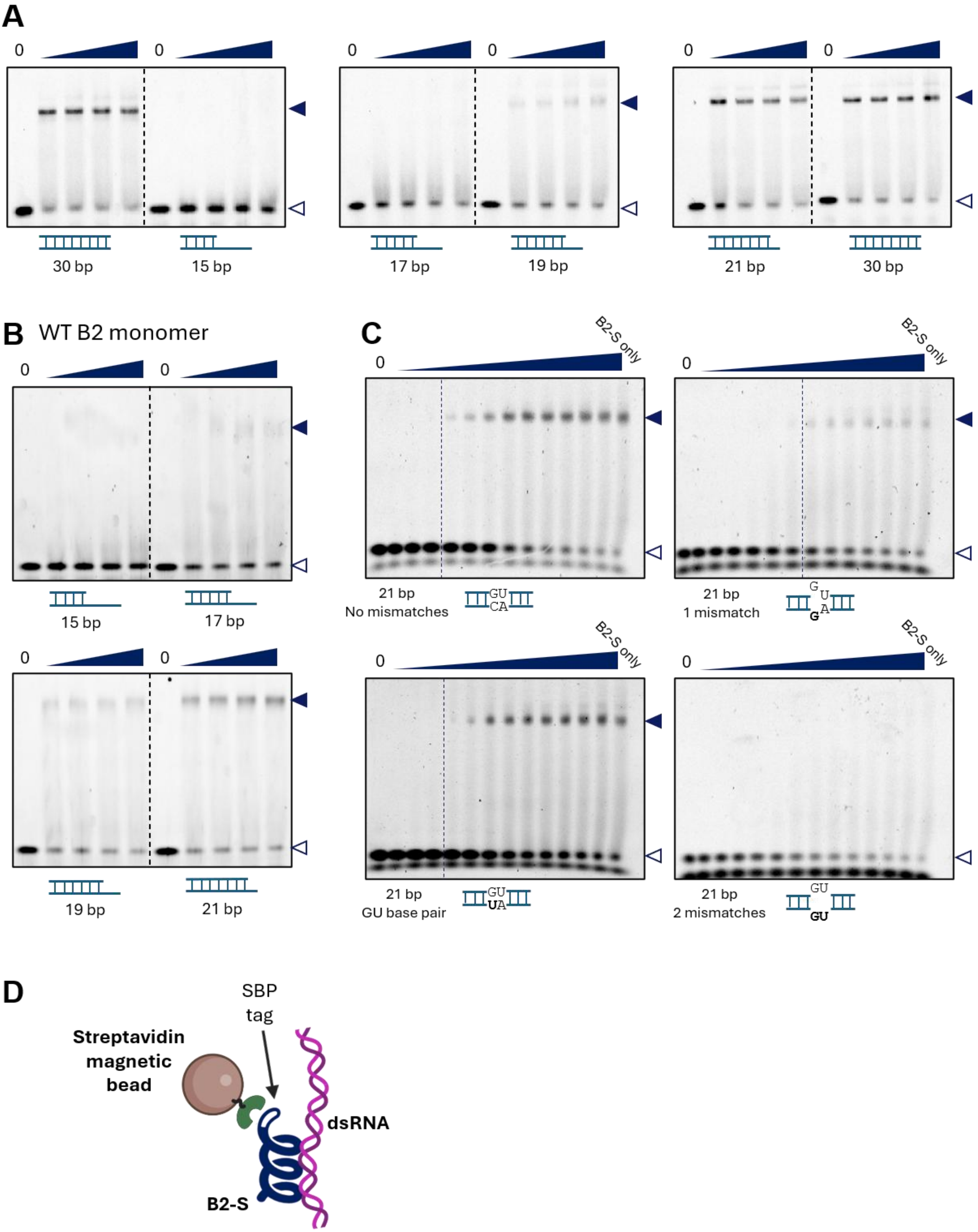
A
0
0
30 bp
15 bp
17 bp
19 bp
21 bp
30 bp
B
WT B2 monomer
15 bp
17 bp
19 bp
21 bp
C
B2-S only
21 bp
No mismatches
GU
CA
21 bp
1 mismatch
G
U
G
A
21 bp
GU base pair
GU
UA
21 bp
2 mismatches
GU
GU
D
SBP
tag
Streptavidin
magnetic
bead
dsRNA
B2-S

**Supplemental Figure 1. B2-S preferentially binds perfectly matched dsRNA substrates that are ≥21 bp. (A and B)** Electrophoretic mobility shift assays (EMSAs) of B2-S (A) or wild-type (WT) B2 monomer (B) with 15-, 17-, 19-, 21-, or 30-bp dsRNA substrates. **(C)** EMSAs of B2-S with a 21-bp dsRNA substrate containing either no mismatch, one mismatch, two mismatches, or a central GU base pair. For each assay in (A–C), a constant amount of dsRNA substrate was incubated with increasing concentrations of B2-S or B2 monomer, as indicated. "B2-S only" lanes contain protein in the absence of dsRNA, whereas "0" lanes contain dsRNA substrate in the absence of protein. Filled arrowheads indicate bound protein–dsRNA complexes, and open arrowheads indicate free dsRNA substrate. **(D)** Schematic illustrating the use of B2-S to capture and remove dsRNA. SBP, streptavidin-binding peptide. Image was made with BioRender.

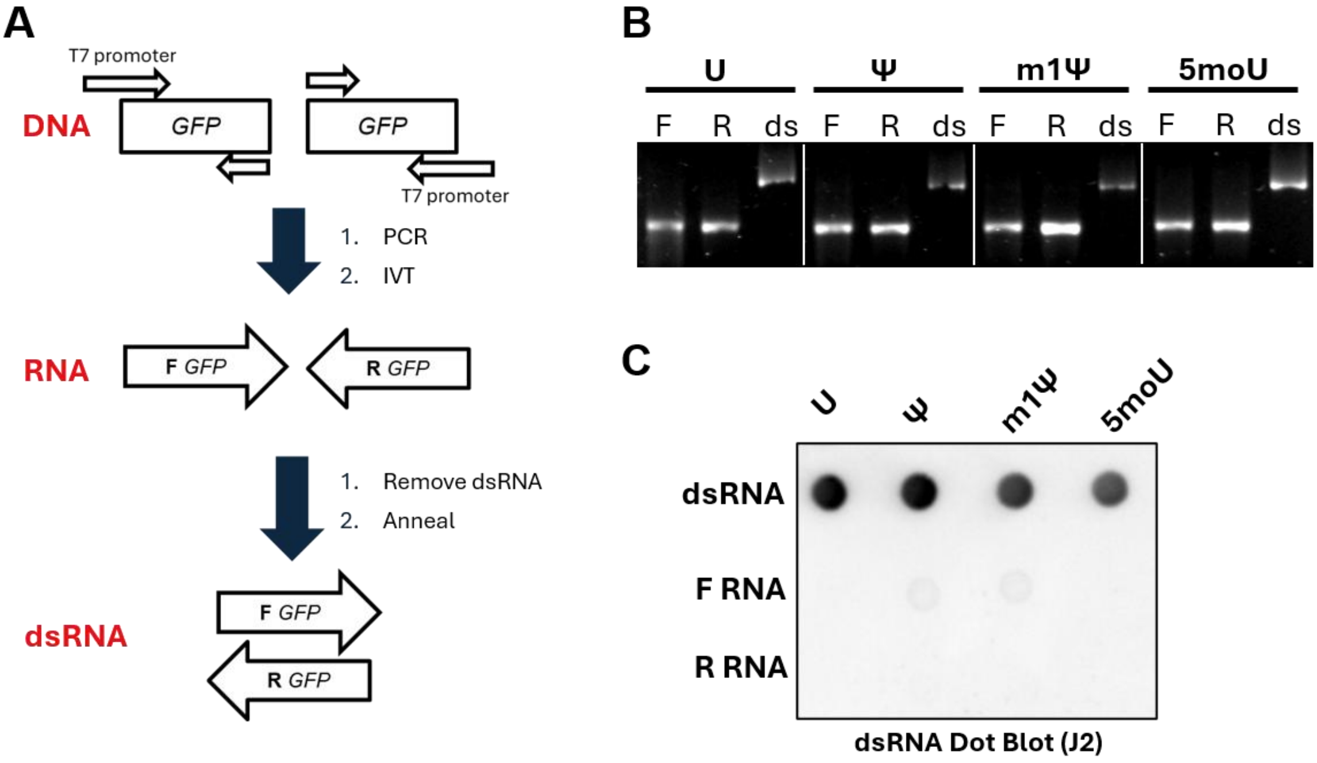


**Supplemental Figure 2. Generation and validation of dsRNA. (A)** Schematic illustrating the workflow used to generate dsRNA. DNA templates containing a T7 promoter were amplified using forward and reverse primers and used for *in vitro* transcription (IVT) to generate complementary forward (F) and reverse (R) RNA strands. Residual dsRNA byproducts were removed from the individual RNA preparations by cellulose–ethanol purification, and the complementary RNAs were subsequently annealed to generate dsRNA. **(B)** Agarose gel analysis of RNA products generated using the indicated modified nucleotides: unmodified U (U), pseudouridine (Ψ), N1-methylpseudouridine (m1Ψ), or 5-methoxyuridine (5moU). F, forward RNA; R, reverse RNA; ds, annealed dsRNA. The slower-migrating band indicates formation of the dsRNA product. **(C)** Dot-blot analysis of the corresponding dsRNA preparations, confirming successful generation of dsRNA containing each indicated uridine modification.

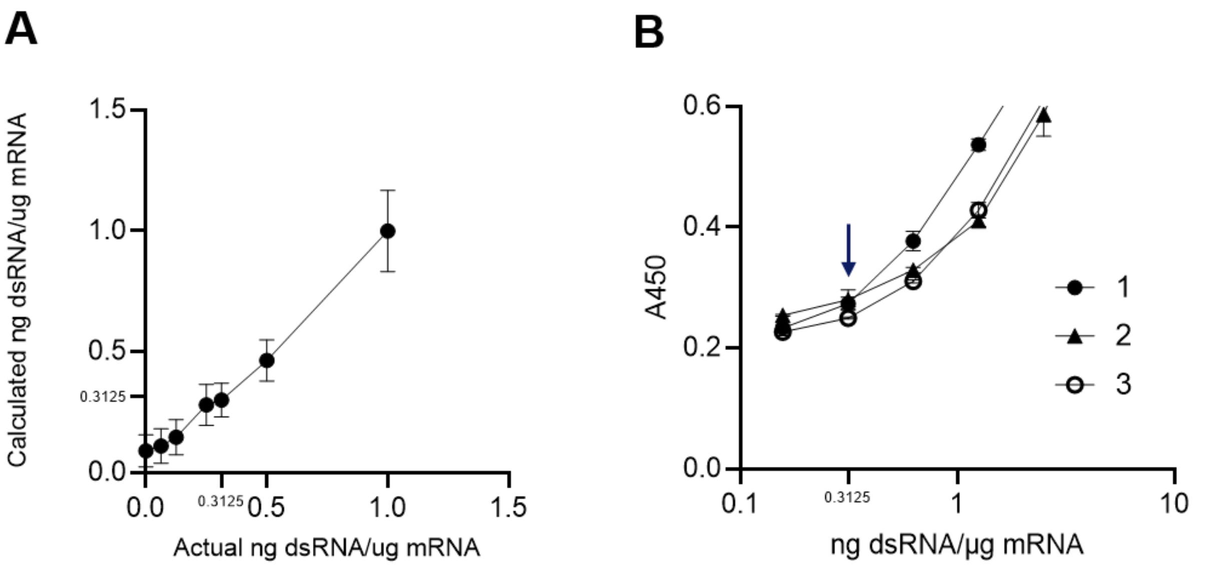


**Supplemental Figure 3. Determination of the lower limit of quantification (LLOQ) for the K1-9D5 ELISA. (A)** HPLC-purified m1Ψ-modified RFP or Cas9 mRNA was spiked with known amounts of m1Ψ-modified dsRNA and analyzed by K1-9D5. K1-9D5 measurements closely matched expected dsRNA concentrations down to 0.3125 ng dsRNA per µg RNA, whereas quantification at lower spike-in levels was increasingly affected by assay background. **(B)** Standard curves from three independent K1-9D5 ELISA runs. The arrow indicates 0.3125 ng dsRNA per µg RNA, corresponding to the transition from the low-signal region to the responsive portion of the curve and defined as the K1-9D5 LLOQ.

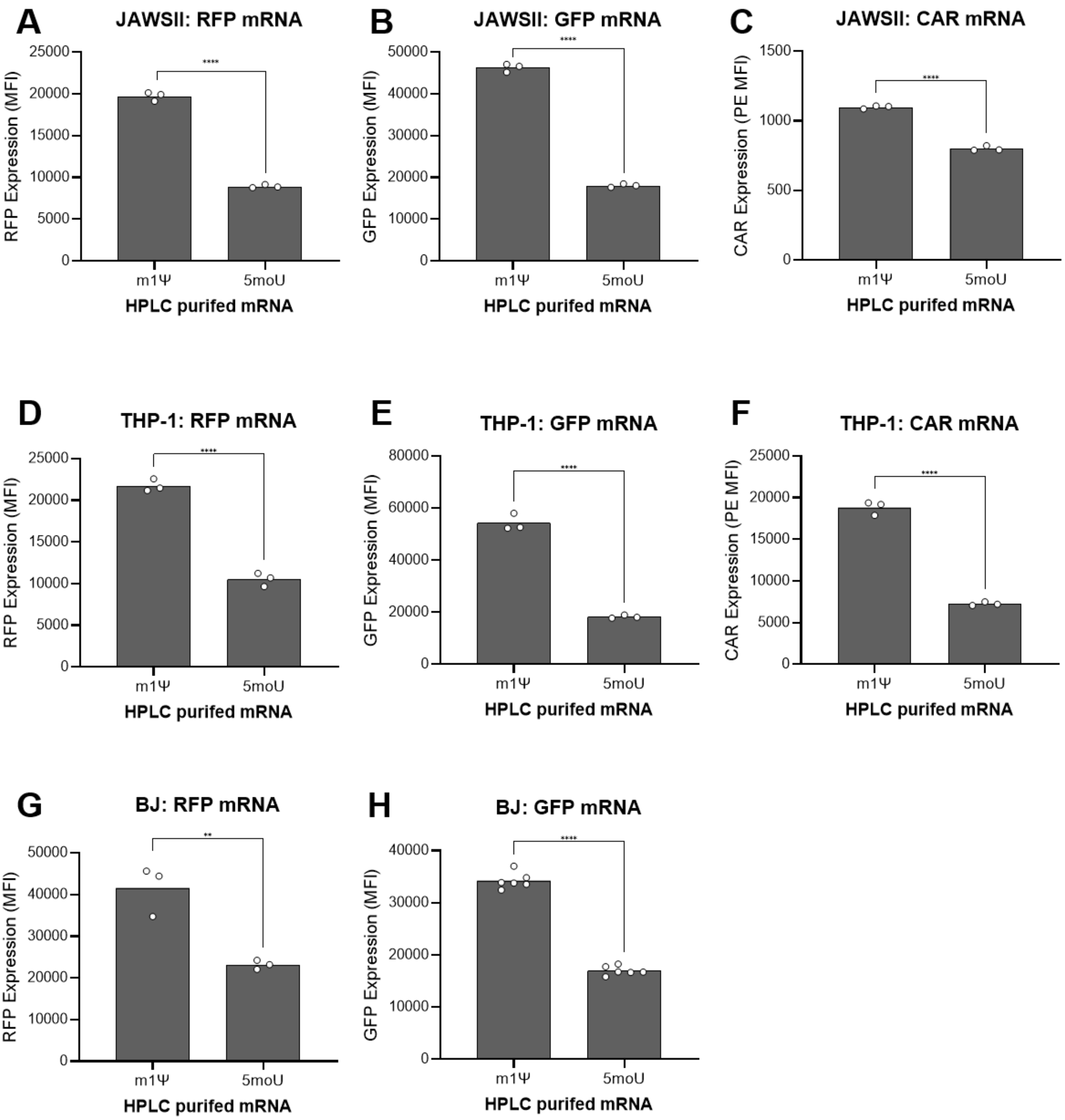


**Supplemental Figure 4. Enhanced expression of HPLC-purified m1Ψ-modified mRNA extends across multiple constructs and cell types.** HPLC-purified mRNAs containing either m1Ψ or 5moU were evaluated for protein expression in multiple cellular contexts. **(A–C)** RFP, GFP, and CD19-CAR expression in murine JAWSII immature dendritic cells. **(D–F)** RFP, GFP, and CD19-CAR expression in human THP-1 monocytes. **(G–H)** RFP and GFP expression in human BJ fibroblast cells. Bars represent the mean fluorescence intensity (MFI) and individual points represent replicates. Statistical significance was determined by unpaired t-test. **: $P \leq 0.01$; ****: $P \leq 0.0001$.

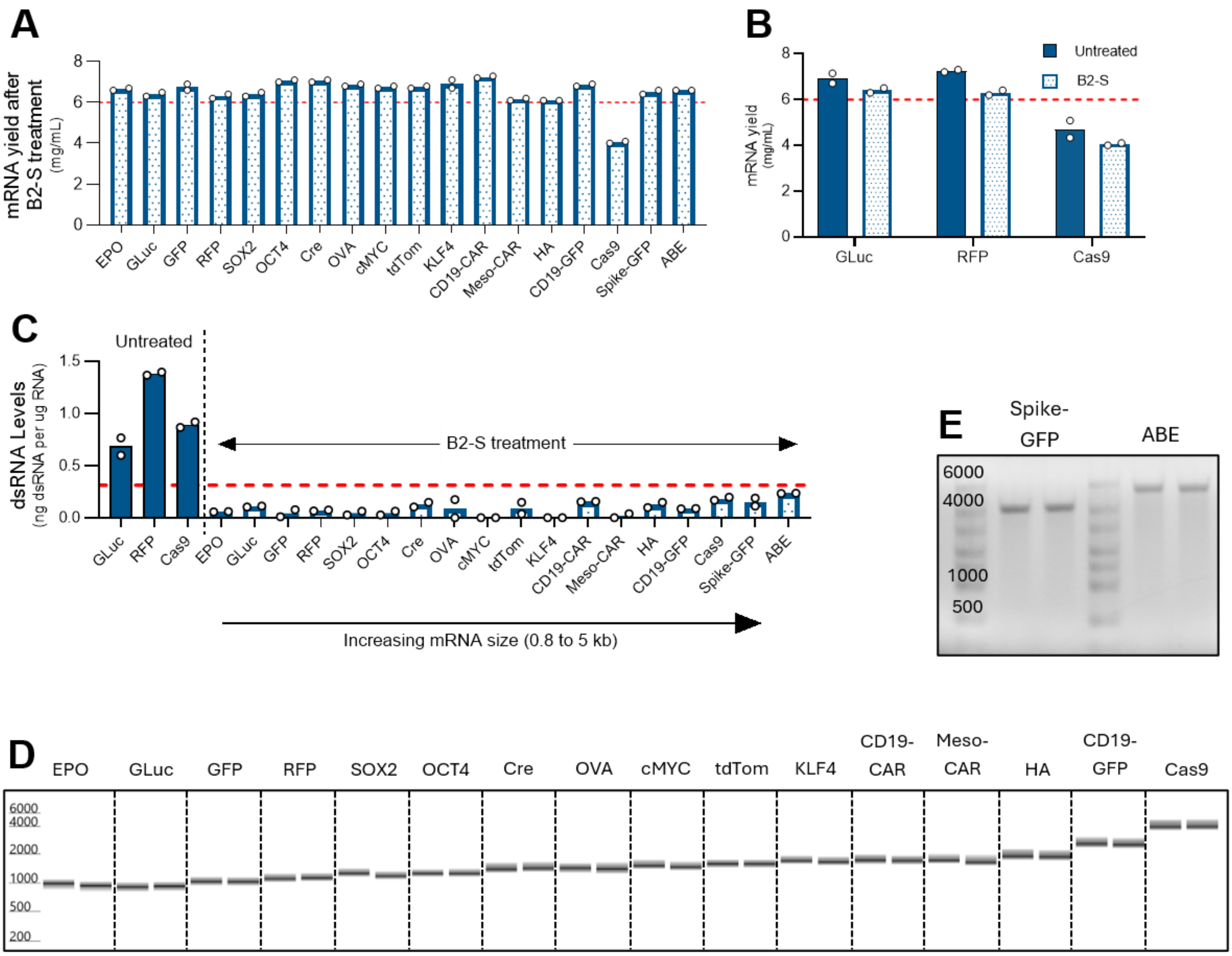


**Supplemental Figure 5. B2-S treatment efficiently removes dsRNA while preserving mRNA yield and integrity across diverse IVT mRNA constructs. (A)** Recovery of IVT mRNA following B2-S treatment across multiple mRNA constructs. The dashed red line indicates an IVT mRNA yield of 6 mg/mL. **(B)** Comparison of mRNA recovery before and after B2-S treatment for selected constructs. The B2-S-treated Cas9 mRNA remained below 6 mg/mL because the corresponding untreated mRNA also had a starting yield below 6 mg/mL, rather than a substantial yield loss during treatment. The dashed red line indicates an IVT mRNA yield of 6 mg/mL. **(C)** dsRNA levels in untreated and B2-S-treated mRNA samples measured by the K1-9D5 ELISA. The dashed red line indicates the lower limit of quantification of 0.3125 ng dsRNA per µg mRNA. B2-S treatment reduced dsRNA to below this threshold across the tested constructs. **(D and E)** Gel-like images from capillary electrophoresis (D) or agarose gel electrophoresis (E) showing mRNA integrity across constructs ranging from approximately 0.8 to 5 kb following B2-S treatment. Bars in plots represent the mean and individual points represent replicate measurements.

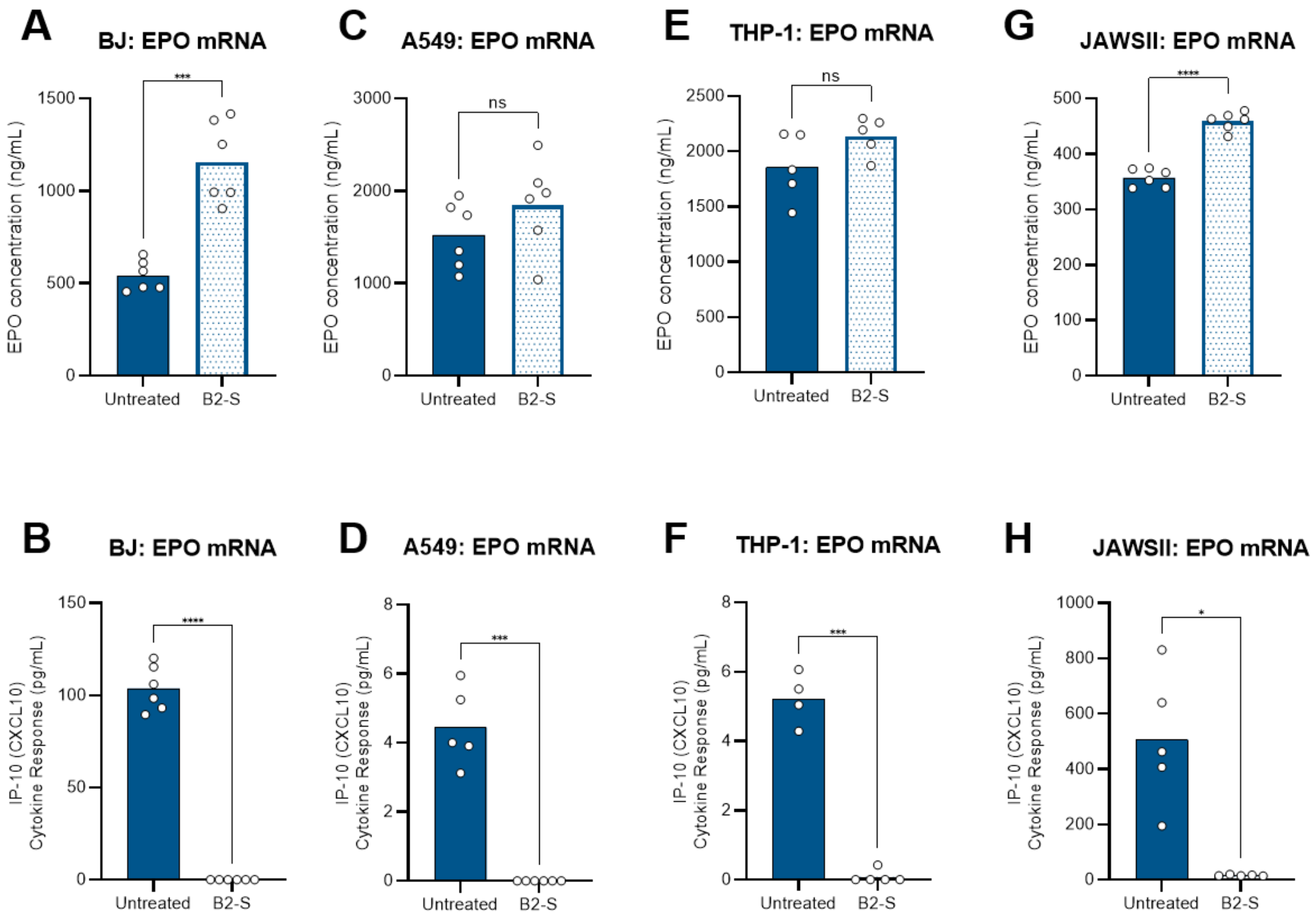


**Supplemental Figure 6. B2-S treatment improves EPO expression while reducing IP-10/CXCL10 induction across multiple cell lines. (Top panels: A, C, E, and G)** EPO expression following transfection with untreated or B2-S-treated EPO mRNA in the indicated cell lines. **(Bottom panels: B, D, F, and H)** IP-10/CXCL10 levels measured in the corresponding samples. B2-S treatment increased EPO expression for most cell lines while markedly reducing IP-10/CXCL10 induction across all the tested cell lines. Bars represent the mean, and individual points represent replicate measurements. Statistical significance was determined by unpaired t-test. *: P≤0.05; ***: P≤0.001; ****: P≤0.0001, **ns**: not significant.

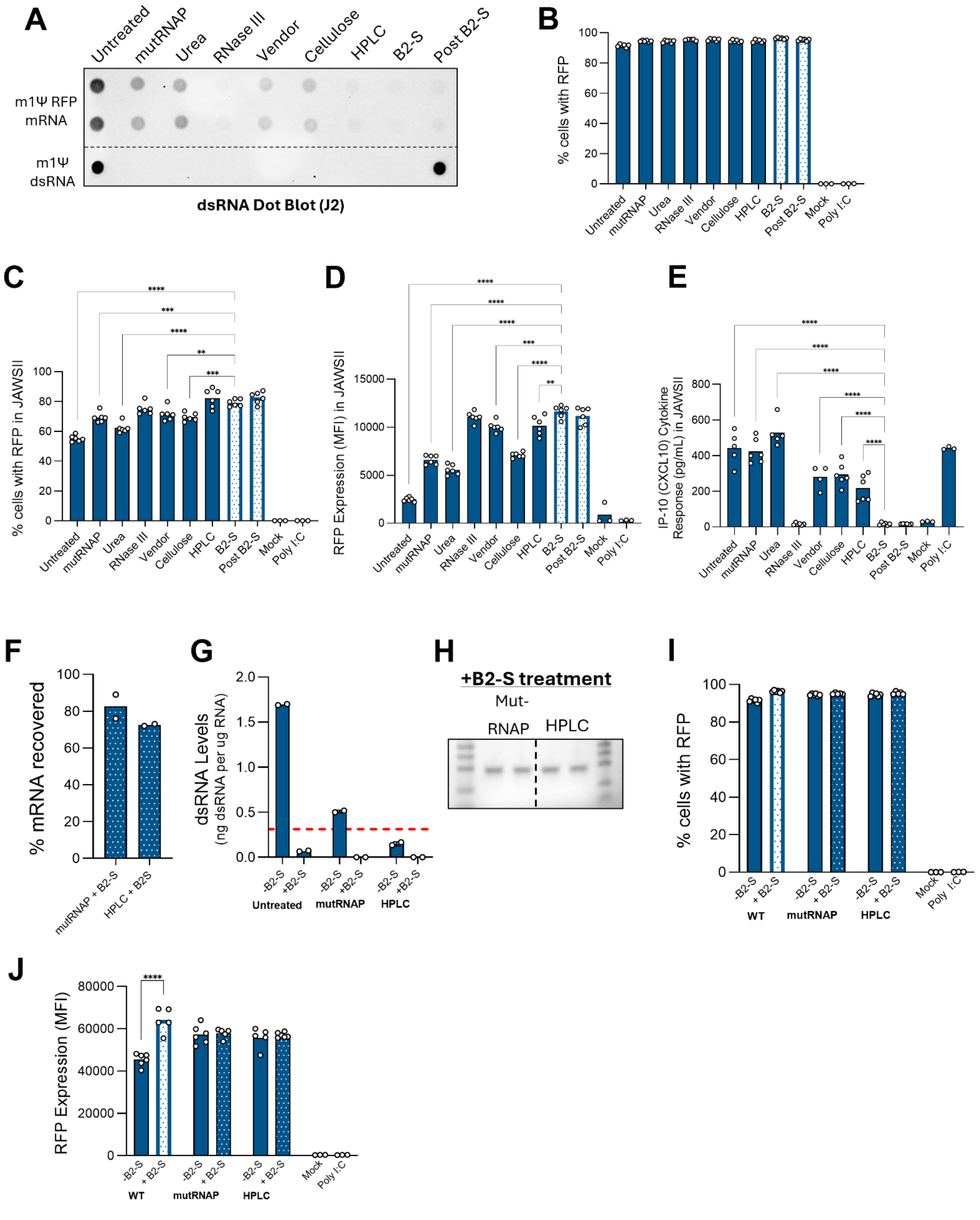
A
Untreated
mutRNAP
Urea
RNase III
Vendor
Cellulose
HPLC
B2-S
Post B2-S
m1Ψ RFP mRNA
m1Ψ dsRNA
dsRNA Dot Blot (J2)
B
% cells with RFP
Mock
Poly I:C
C
% cells with RFP in JAWSII
D
RFP Expression (MFI) in JAWSII
E
IP-10 (CXCL10) Cytokine Response (pg/mL) in JAWSII
F
% mRNA recovered
mutRNAP + B2-S
HPLC + B2S
G
dsRNA Levels (ng dsRNA per ug RNA)
-B2-S
+B2-S
Untreated
mutRNAP
HPLC
H
+B2-S treatment
Mut-RNAP
HPLC
I
% cells with RFP
WT
J
RFP Expression (MFI)

**Supplemental Figure 7. Functional and analytical comparison of dsRNA-reduction workflows and post-hoc B2-S treatment. (A)** J2 dot blot analysis of residual dsRNA in RFP mRNA prepared using the indicated dsRNA-reduction workflows. **(B)** Percentage of RFP-positive BJ fibroblasts 24 h after transfection with RFP mRNA generated using the indicated workflows. **(C–E)** Functional evaluation of the same RFP mRNA preparations in JAWSII cells, including the percentage of RFP-positive cells (C), RFP expression (D), and IP-10/CXCL10 secretion (E). **(F–J)** Characterization of mutant RNAP- and HPLC-purified RFP mRNA before and after post-hoc B2-S treatment. mRNA recovery (F), residual dsRNA measured by K1-9D5 ELISA (G), mRNA integrity (H), percentage of RFP-positive cells (I), and RFP expression (J) were evaluated in BJ fibroblasts following B2-S treatment. The dashed red line in (G) indicates the K1-9D5 lower limit of quantification (LLOQ) of 0.3125 ng dsRNA per µg mRNA. Mock-transfected and poly(I:C)-treated cells were included as negative and positive controls, respectively, where indicated. Bars represent the mean, and individual replicate values are shown as open circles. Statistical significance was determined in (C-E) by one-way ANOVA followed by Dunnett's multiple comparisons test versus the B2-S sample and in (I-J) by two-way ANOVA followed by Tukey’s multiple comparisons test versus the respective B2-S-treated sample. Not significant comparisons are not indicated. **: $P \leq 0.01$; ***: $P \leq 0.001$; ****: $P \leq 0.0001$.

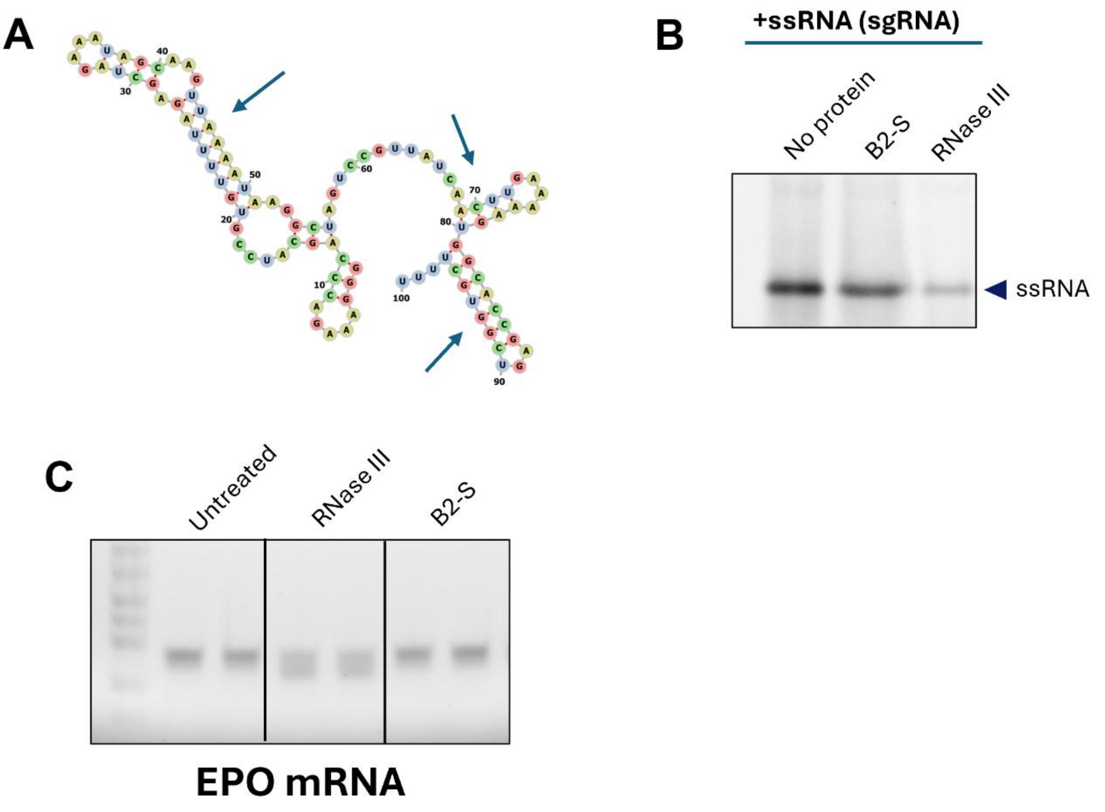


**Supplemental Figure 8. RNase III sensitivity is substrate dependent, whereas B2-S treatment preserves RNA integrity. (A)** Predicted secondary structure of the sgRNA used in this study. See Supplemental Methods for the sequence. Nucleotides are color coded by base identity: adenine (A), yellow; uracil (U), blue; guanine (G), red; and cytosine (C), green. Arrows indicate intramolecular base-paired regions with dsRNA-like structure. Image was generated with RNAfold (44). **(B)** Gel analysis of sgRNA following the indicated treatments, showing degradation after RNase III treatment compared with preservation of the intact RNA species following B2-S treatment. **(C)** Gel analysis of EPO mRNA following the indicated treatments, demonstrating RNase III-associated degradation and preservation of mRNA integrity following B2-S treatment.

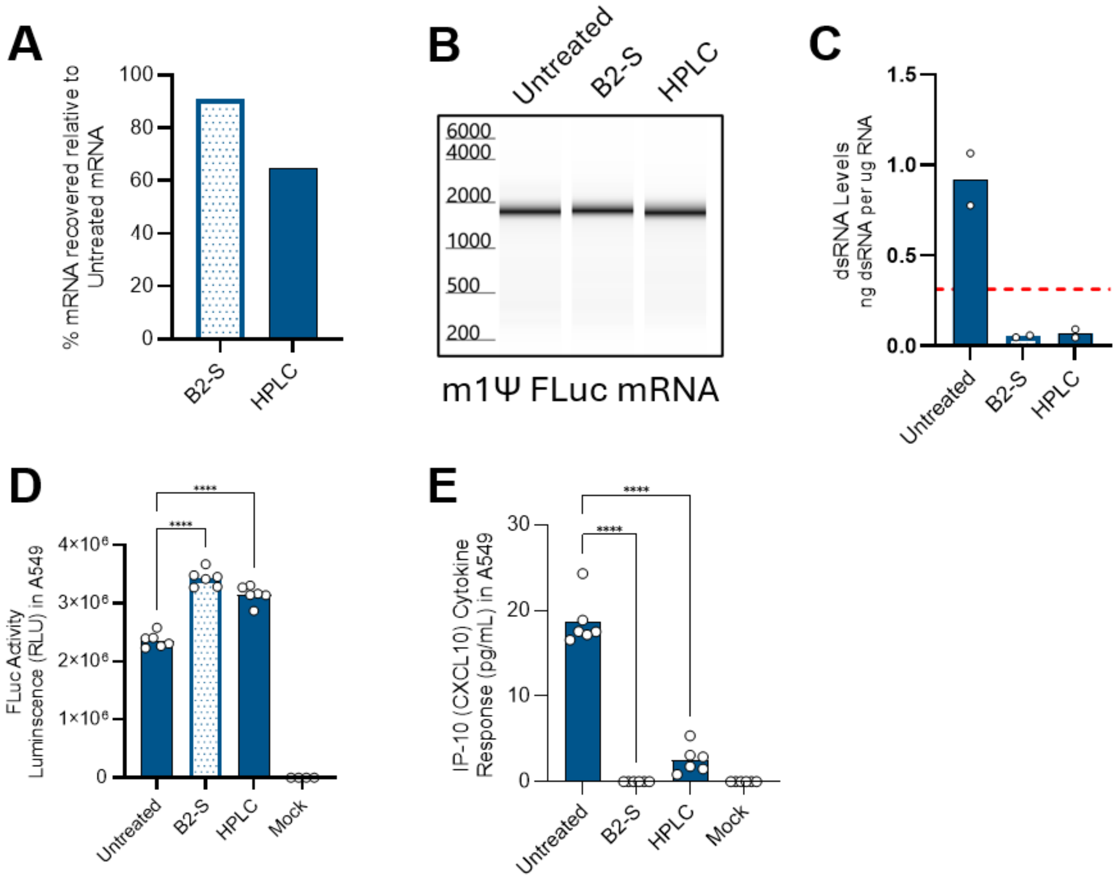


**Supplemental Figure 9. B2-S treatment preserves FLuc mRNA recovery and integrity while reducing dsRNA and improving functional performance. (A)** Recovery of m1Ψ-modified firefly luciferase (FLuc) mRNA following B2-S treatment or HPLC purification relative to untreated mRNA. **(B)** Capillary electrophoresis analysis showing preservation of FLuc mRNA integrity following B2-S treatment and HPLC purification. **(C)** Residual dsRNA levels in untreated, B2-S-treated, and HPLC-purified FLuc mRNA measured by the K1-9D5 ELISA. The dashed red line indicates the assay LLOQ. **(D)** FLuc activity in A549 cells following transfection with untreated, B2-S-treated, or HPLC-purified FLuc mRNA. **(E)** IP-10/CXCL10 levels in culture media from the corresponding A549 cell samples. Bars represent the mean, and individual replicate values are shown as open circles. Statistical significance was determined by one-way ANOVA followed by Tukey's multiple comparisons test versus the B2-S sample. Not significant comparisons are not indicated. ********: P≤0.0001.

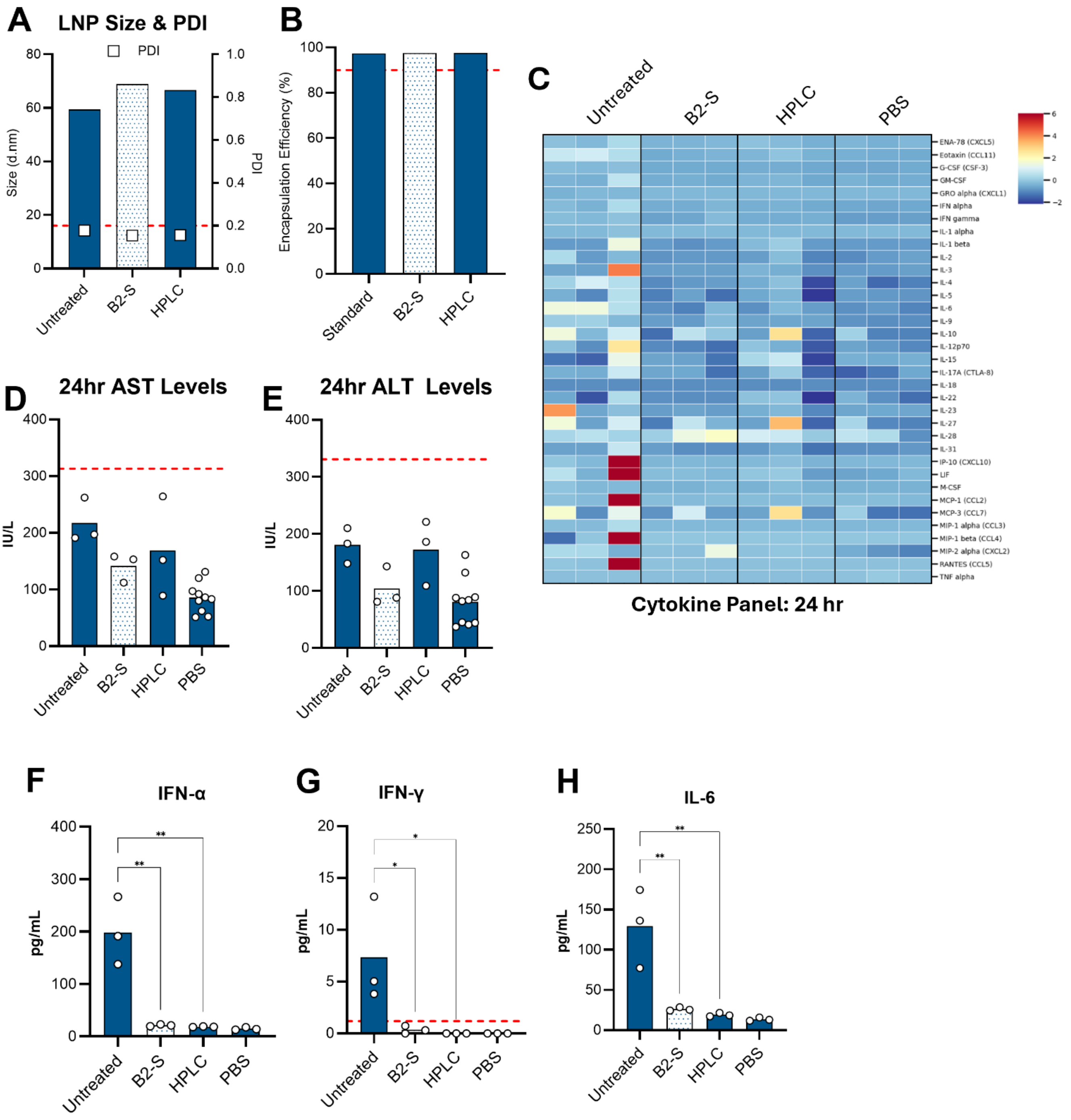


**Supplemental Figure 10. B2-S-treated mRNA shows low *in vivo* toxicity and inflammatory signaling. (A)** Particle size and polydispersity index (PDI) of LNPs formulated with untreated, B2-S-treated, or HPLC-purified FLuc mRNA. Particle size is plotted on the left y-axis and PDI on the right y-axis. **(B)** Encapsulation efficiency of FLuc mRNA in the corresponding LNP formulations. The red dotted line indicates 90%. **(C)** Serum cytokine and chemokine profiles measured by 36-plex ProcartaPlex at 24 h after intravenous administration of FLuc mRNA-LNPs. Values are displayed as Z-scores (pg/mL) for each analyte across treatment groups. **(D and E)** Serum aspartate aminotransferase (AST) (D) and alanine aminotransferase (ALT) (E) levels following LNP administration. Dashed red lines indicate three times the upper limit of normal (3× ULN). **(F–H)** Serum IFN-α (F), IFN-γ (G), and IL-6 (H) levels 4 hours following administration of untreated, B2-S-treated, or HPLC-purified FLuc mRNA-LNPs. The dashed red line in (G) indicates the lower limit of detection (LLOD) for IFN-γ of 1.2 pg/mL. PBS-treated animals are included as controls where indicated. Bars represent the mean, and individual animals are shown as open circles. Statistical significance was determined by one-way ANOVA followed by Tukey's multiple comparisons test. Not significant comparisons are not indicated. *****: P≤0.05; ******: P≤0.01.

**sgRNA sequence for RNAfold**
5'-GGGAAAGACCCAGCAUCCGUGUUUUAGAGCUAGAAAUAGCAAGUUAAAAUAAGGCUAGUC
CGUUAUCAACUUGAAAAAGUGGCACCGAGUCGGUGCUUUUU-3'

The underlined portion is the gRNA spacer.
